# Multi-omic spatial profiling reveals the unique virus-driven immune landscape of COVID-19 placentitis

**DOI:** 10.1101/2022.11.14.516398

**Authors:** Matthew Pugh, Éanna Fennell, Ciara I Leahy, Tracey Perry, Beata Hargitai, Tamas Marton, Kelly J. Hunter, Graham Halford, Hale Onder Yilmaz, Zania Stamataki, Gary Reynolds, Harriet J. Hill, Benjamin E. Willcox, Neil Steven, Catherine A. Thornton, Stefan D. Dojcinov, Aedin Culhane, Paul G. Murray, Graham S. Taylor

**Author notes:** These authors contributed equally (Matthew Pugh, Éanna Fennell). Corresponding Authors (Correspondence &).

## Abstract

COVID-19 placentitis, a rare complication of maternal SARS-CoV-2 infection, only shows detectable virus in the placenta of a subset of cases. We provide a deep multi-omic spatial characterisation of placentitis from obstetrically complicated maternal COVID-19 infection. We found that SARS-CoV-2 infected placentas have a distinct transcriptional and immunopathological signature. This signature overlaps with virus-negative cases supporting a common viral aetiology. An inverse correlation between viral load and disease duration suggests viral clearance over time. Quantitative spatial analyses revealed a unique microenvironment surrounding virus-infected trophoblasts characterised by PDL1-expressing macrophages, T-cell exclusion, and interferon blunting. In contrast to uninfected mothers, ACE2 was localised to the maternal side of the placental trophoblast layer of almost all mothers with placental SARS-CoV-2 infection, which may explain variable susceptibility to placental infection. Our results demonstrate a pivotal role for direct placental SARS-CoV-2 infection in driving the unique immunopathology of COVID-19 placentitis.

**Graphical Abstract:** 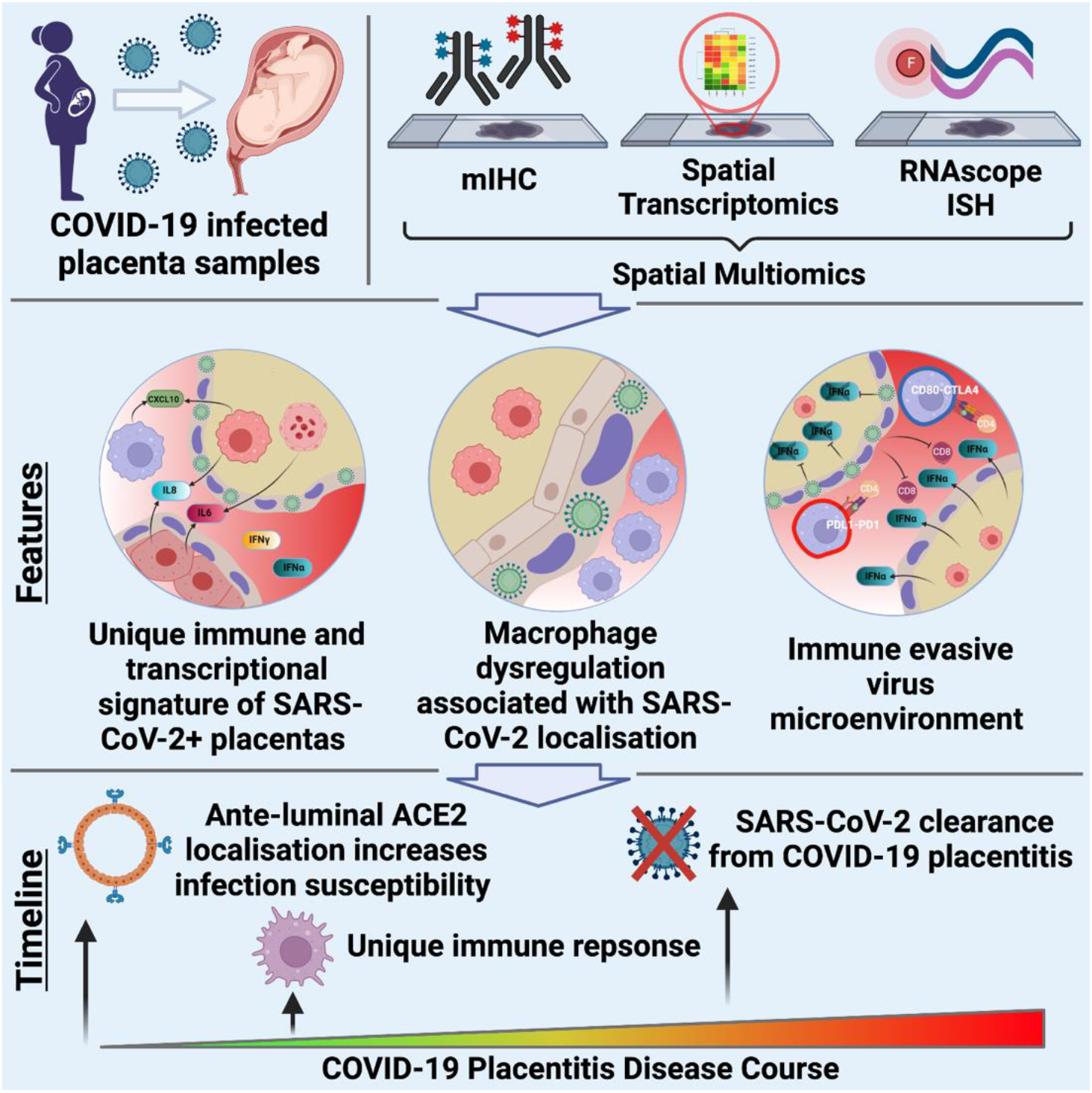

## Introduction

The placenta forms an anatomical barrier between the maternal and fetal blood supplies^1^. Placental integrity is crucial to protect the fetus from infection and to prevent immune rejection of the semi-allogeneic fetus. The placenta predominantly comprises fetal-derived villi lined by a bilayer of cytotrophoblasts and fused syncytiotrophoblasts, collectively referred to as the trophoblast layer. The villi, which contain circulating fetal blood, invade the maternal uterine decidua and are surrounded by the intervillous space through which maternal blood flows allowing nutrient and gas exchange to occur. Diverse mechanisms operate at the maternal-fetal interface to prevent or limit virus infections^2^ including: constitutive and inducible interferon expression; proinflammatory cytokine responses; and the presence of maternal-derived innate cells in the decidua and fetal-derived macrophages, called Hofbauer cells, in the villous stroma^3^.

Several viruses, such as Zika and cytomegalovirus, have the potential to cause harm to the fetus, either through the systemic effects of virus infection or, despite the placenta’s anti-viral defences, through direct infection of the placenta^4,5^. Recently, it has been shown that severe respiratory COVID-19 during pregnancy is associated with an increased risk of serious obstetric complications which can include intrauterine growth restriction, premature labour and stillbirth^6,7^, raising the possibility that these adverse outcomes could be due to systemic effects of infection^8^. However, severe obstetric complications associated with characteristic inflammation of the placenta, termed COVID-19 placentitis, have also been reported in women with mild or asymptomatic respiratory infection ^9–11^. COVID-19 placentitis has three characteristic pathological features: macrophage infiltration in the intervillous space, termed chronic histiocytic intervillositis (CHI); fibrin deposition in the intervillous space amounting to massive perivillous fibrin deposition (MPVFD); and extensive trophoblast necrosis (TN)^12^. These pathologies typically affect most of the placenta, leading to placental destruction and fetal demise in most cases^13^.

The exact role of SARS-CoV-2 in COVID-19 placentitis is unclear. For example, MPVFD and CHI occur in other placental pathologies that predate the pandemic, raising the possibility that the virus is opportunistically infecting compromised placental tissues rather than being an initiating event. Moreover, because placental SARS-CoV-2 is detected in only one-half of COVID19 placentitis cases it has been suggested that systemic inflammation rather than being direct placental infection is the pathogenic driver of disease^10,14,15^. However, the occurrence of severe obstetric complications and placentitis in mothers with mild or asymptomatic respiratory COVID-19 argues against a dominant role for systemic inflammation^16^. This raises the alternative possibility that direct placental infection is centrally important but is subsequently cleared, as has recently been reported for SARS-CoV-2 infection of the lungs^17,18^.

To define the role of the virus in the pathogenesis of COVID-19 placentitis we integrated bulk and spatially resolved tissue transcriptomics with quantitative high-dimensional immunohistochemistry to comprehensively characterise the immunopathology of this disease. Collectively, our data identify a unique virus associated immunopathological signature, features of which are shared with temporally more advanced cases lacking detectable virus, indicating viral loss. Our observations shed new insights into the biology of placentitis, reveal potential novel therapeutic targets and point to the need for a re-evaluation of the diagnostic criteria used to define COVID-19 placentitis. Our study also suggests that the impact of SARS-CoV-2 in mediating obstetric complications is likely to be greater than previously appreciated.

## Results

### Study cohort, sample collection and multi-omic analysis

Formalin fixed paraffin embedded (FFPE) placental tissues from 13 SARS-CoV-2 infected mothers suffering adverse obstetric outcomes were obtained from a UK tertiary regional centre (Figure 1A). One case had previously been published in a case report and 6 cases were published as part of a case series^11,16^. Respiratory SARS-CoV-2 infection was detected by PCR test following routine nasopharyngeal swab in 12/13 (92%) cases. One mother tested negative by respiratory PCR but was subsequently identified as virus-positive based on SARS-CoV-2 detection in her placenta. Eleven pre-pandemic un-infected control placentas, comprising 4 CHI controls (Control^CHI^), 2 chronic villitis (CV) controls (Control^CV^) and 5 normal controls (Normal), were collected from the same centre.

**Figure 1.**
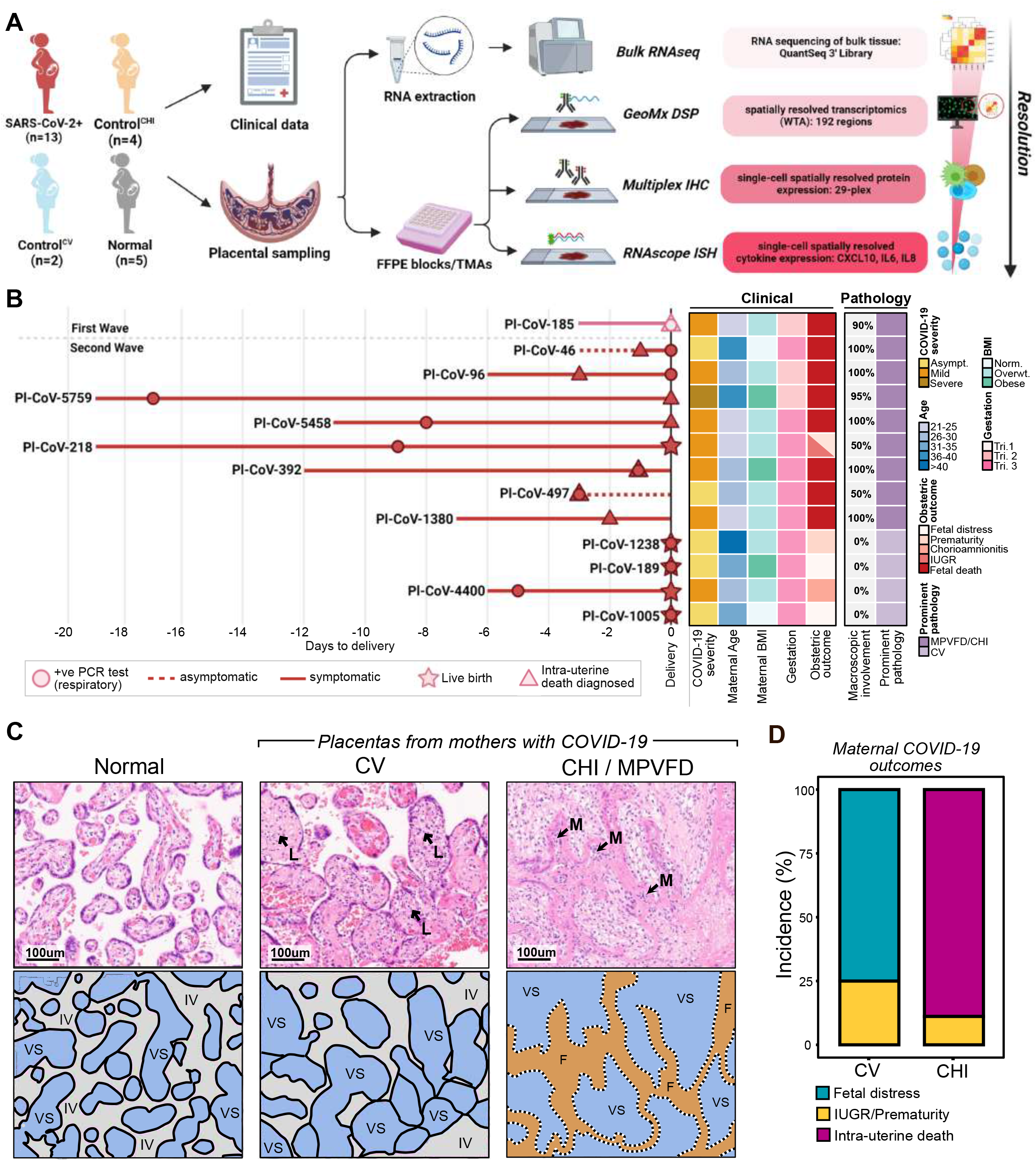
Study overview and clinicopathological features of the cohort. (A) Overview illustration showing the multi-omic workflow of placental samples from 13 SARS-CoV-2 infected women and 11 uninfected controls. (B) Timeline of clinical events relative to time of delivery. Heatmap displays clinical and pathological features for each SARS-CoV-2 infected mother. (C) Examples of the two morpho-pathological categories of placentas from SARS-CoV-2 infected mothers: chronic histiocytic intervillositis with massive perivillous fibrin deposition (CHI/MPVFD) and chronic villitis (CV). Normal placental pathology is also included as a control. IV - Intervillous space; VS – Villous stroma; F – Fibrin; L – Lymphocytes; M – Macrophages. (D) Obstetric outcome of SARS-CoV-2 infected mothers stratified on morpho-pathological category. See also Figure S1 and Table S1

Following routine clinico-pathological assessment, placenta samples were analysed by four complementary multi-omic approaches: i) Tissue-wide whole-transcriptome analysis using bulk Quantseq 3’ RNA sequencing; ii) Spatially-resolved whole-transcriptome analysis by NanoString GeoMx digital spatial profiling (DSP), targeting the trophoblast, immune, stromal and decidual compartments; iii) Targeted cellular-level expression of specific viral and cytokine genes by multiplex RNAscope in conjunction with TSA-based immunohistochemistry, and; iv) spatial tissue proteomics using multiplex immunohistochemistry (mIHC) on the Lunaphore COMET platform. The latter employed a validated panel of 29 antibodies for detection of cell identity and function (including immune checkpoint expression) as well as expression of virus receptor (ACE2) and viral spike and nucleocapsid proteins (Figure 1A and S1).

### Histological features in placentas from SARS-CoV-2 infected mothers correlate with obstetric outcome

The clinical characteristics of the cohort are described in Figure 1B and supplementary Table 1. Notably, most SARS-CoV-2 infected mothers were either asymptomatic (5/13 - 38%) or had mild respiratory illness (7/13 - 54%); only 1/13 (8%) had severe respiratory illness requiring intensive therapy unit (ITU) admission. The obstetric outcomes, however, were severe. Intrauterine death was the most common adverse obstetric outcome (8/13 - 62%), followed by fetal distress (3/13 - 23%) and combined premature labour with intrauterine growth restriction (2/13 - 15%).

On pathological assessment, 9/13 (69%) of the placentas from SARS-CoV-2 infected mothers showed widespread (>50%) macroscopic lesional involvement (Figure 1B and S1) and histological features consistent with COVID-19 placentitis: CHI, with MPVFD and extensive trophoblast necrosis - hereafter denoted as CoV19^CHI 16^. Compared to pre-pandemic control^CHI^ cases, these nine CoV19^CHI^ cases showed much more extensive trophoblast necrosis and, unlike control^CHI^, this often involved the circumference of the placental villi (Figure 1C and S1). Intervillous histiocytes in the CoV19^CHI^ cases also showed a distinctive distribution, abutting placental trophoblasts, resulting in histiocytic palisading (Figure S1), a feature not observed in the control^CHI^ cases. The remaining 4/13 (31%) cases from SARS-CoV-2 infected mothers showed no evidence of macroscopic disease and histologically had a chronic villitis (CV) predominant pattern of inflammation, hereafter denoted as CoV19^CV^. There was a significant difference in obstetric outcomes between CoV19^CHI^ and CoV19^CV^ with 8/9 (89%) cases of the former resulting in intrauterine death compared to 0/4 (0%) of the latter (p=0.007 Fisher exact test, Figure 1D).

### SARS-CoV-2 infects and replicates in placental trophoblasts of a subset of cases of COVID-19 placentitis

Placental mIHC data from all 13 SARS-CoV-2 infected mothers and 6 controls were initially assessed for virus positivity. To ensure robust virus detection, positive cases were defined only as those which showed co-expression of spike and nucleocapsid proteins using antibodies we validated for specificity (Figure S2). SARS-CoV-2 was detected in 6/13 (46%) placentas from SARS-CoV-2 infected mothers and in none of the controls. All six virus-positive placentas were CoV19^CHI^ cases (virus positive and negative cases denoted as P^+^CoV19 and P^-^CoV19, respectively, Figure 2A). Virus proteins were not detectable in the four CoV19^CV^ cases. To confirm virus status, we aligned mRNA transcripts from the bulk RNAseq data to the SARS-CoV-2 genome and performed RNAscope *in-situ* hybridization (ISH) using viral ORF1ab sense and spike antisense probes (to detect viral transcription and genome, respectively). Of the 6 placentas positive for virus protein by mIHC, 5 were also tested by RNA analysis and all were positive (Figure 2B - E).

**Figure 2.**
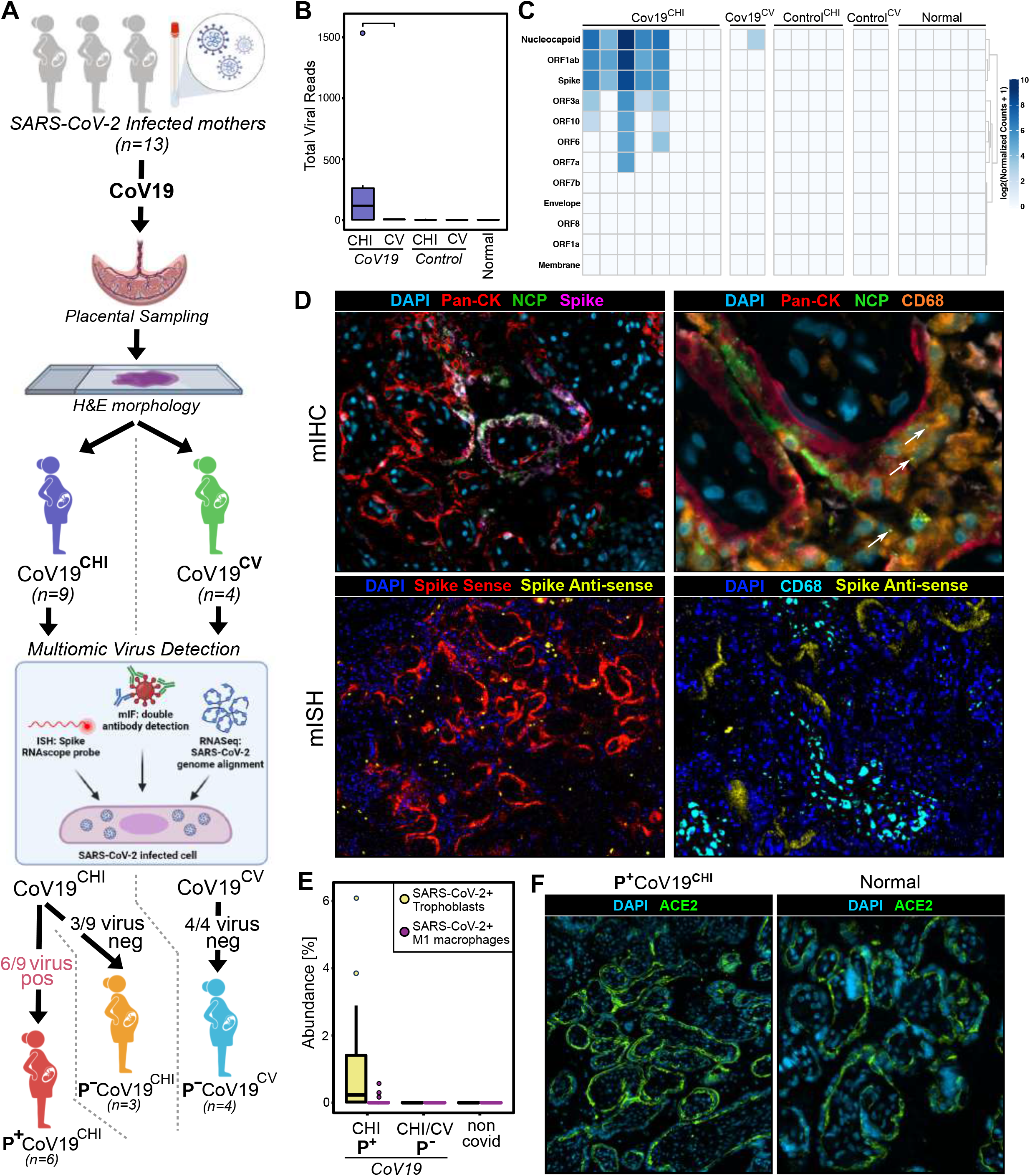
Multi-omic virus detection in placentas from SARS-CoV-2 positive mothers. (A) Multi-omic cross-validated detection of SARS-CoV-2 and case stratification based on viral status and morpho-pathological features of the placenta. (B and C) Bulk RNAseq analysis of placenta showing (B) total viral reads and (C) individual viral genes. (D) Virus localisation within placenta by multiplex immunofluorescence (mIHC, top panels). and RNAscope multiplex in situ hybridisation (mISH, bottom panels). Arrows in the mIHC images indicate viral nucleocapsid positive granules in CD68-positive macrophages surrounding virus infected trophoblasts. (E) Abundance of SARS-CoV-2 positive trophoblasts and macrophages in placenta tissue determined by mIHC. (F) ACE2 protein detection in placental trophoblasts in SARS-CoV-2 infected tissue and normal uninfected controls determined by mIHC. See also Figure S2.

In the virus-positive placentas, bulk RNAseq showed nucleocapsid, spike and ORF1ab were the most consistent and abundantly expressed virus genes (Figure 2C). Both mIHC and ISH showed that SARS-CoV-2 was located predominantly in the syncytiotrophoblast layer with a patchy geographical distribution (Figure 2D). ISH also showed these cells contained antisense ORF1ab transcripts indicating replicative infection. We also observed SARS-CoV-2 spike and nucleocapsid proteins in a small population of intervillous macrophages abutting virus-infected trophoblasts (Figure 2D, E). However, in contrast to infected syncytiotrophoblasts, the viral proteins in these macrophages had a granular cytoplasmic distribution consistent with their localisation to phagocytic vacuoles (Figure 2D). We did not detect viral RNA in macrophages. RT-PCR and Sanger sequencing of the spike gene from two P^+^CoV19^CHI^ placental samples, collected in the first and second UK pandemic waves, showed the virus strain in the placenta reflected the local circulating strains in the community and that both ancestral hCoV-19/Wuhan/WIV04 and the alpha variant can infect the placenta (Figure S2).

### Syncytiotrophoblasts from virus-infected mothers uniquely express ACE2 on the maternal luminal surface

Differences in trophoblast susceptibility to virus entry might explain the absence of SARS-CoV-2 in the placentas of some infected mothers. We measured expression of the virus receptor *ACE2* and of other genes involved in virus entry, including *BSG1, NRP1* and *TMPRSS2*, in the trophoblast layer using NanoString GeoMx digital spatial profiling. All four genes were expressed in all sub-groups of placentas, irrespective of virus presence, with no significant upregulation of expression in regions enriched for infected trophoblasts (Figure S2). However, while mIHC revealed that all placental subgroups showed equivalent levels of ACE2 protein in the trophoblast layer it also identified a marked difference in ACE2 distribution on the trophoblasts of SARS-CoV-2 infected mothers (Figure 2F). Thus, in controls, including pre-pandemic CHI, ACE2 was confined to the fetal ante-luminal surface, whereas in the diseased placentas of 12/13 SARS-CoV-2 infected mothers, irrespective of placental viral status, ACE2 was expressed in a ‘tramtrack’ distribution on both fetal ante-luminal and maternal luminal surfaces of trophoblasts. Thus, ACE2 on the trophoblast layer is exposed to the maternal circulation in virus-infected mothers but not in placentas from uninfected mothers including those with CHI or CV.

### SARS-CoV-2 infected placentas exhibit a unique transcriptional and immune signature

To further characterise the immune landscape of placentas from SARS-CoV-2 infected mothers, the COMET mIHC data was re-assessed and validated by three pathologists (MP, TM, BH). Consistent with the H&E morphology, all CoV19^CHI^ placentas, regardless of virus status, showed extensive perivillous fibrin deposition and a mixed immune intervillous infiltrate (Figure 3A). The infiltrate was comprised of CD68^+^ macrophages with smaller populations of CD20^+^ B-cells, CD3^+^CD4^+^ and CD3^+^CD8^+^ T-cells, and CD15^+^ neutrophils. In contrast, the intervillous infiltrate of pre-pandemic control^CHI^ consisted almost entirely of CD68+ macrophages.

**Figure 3.**
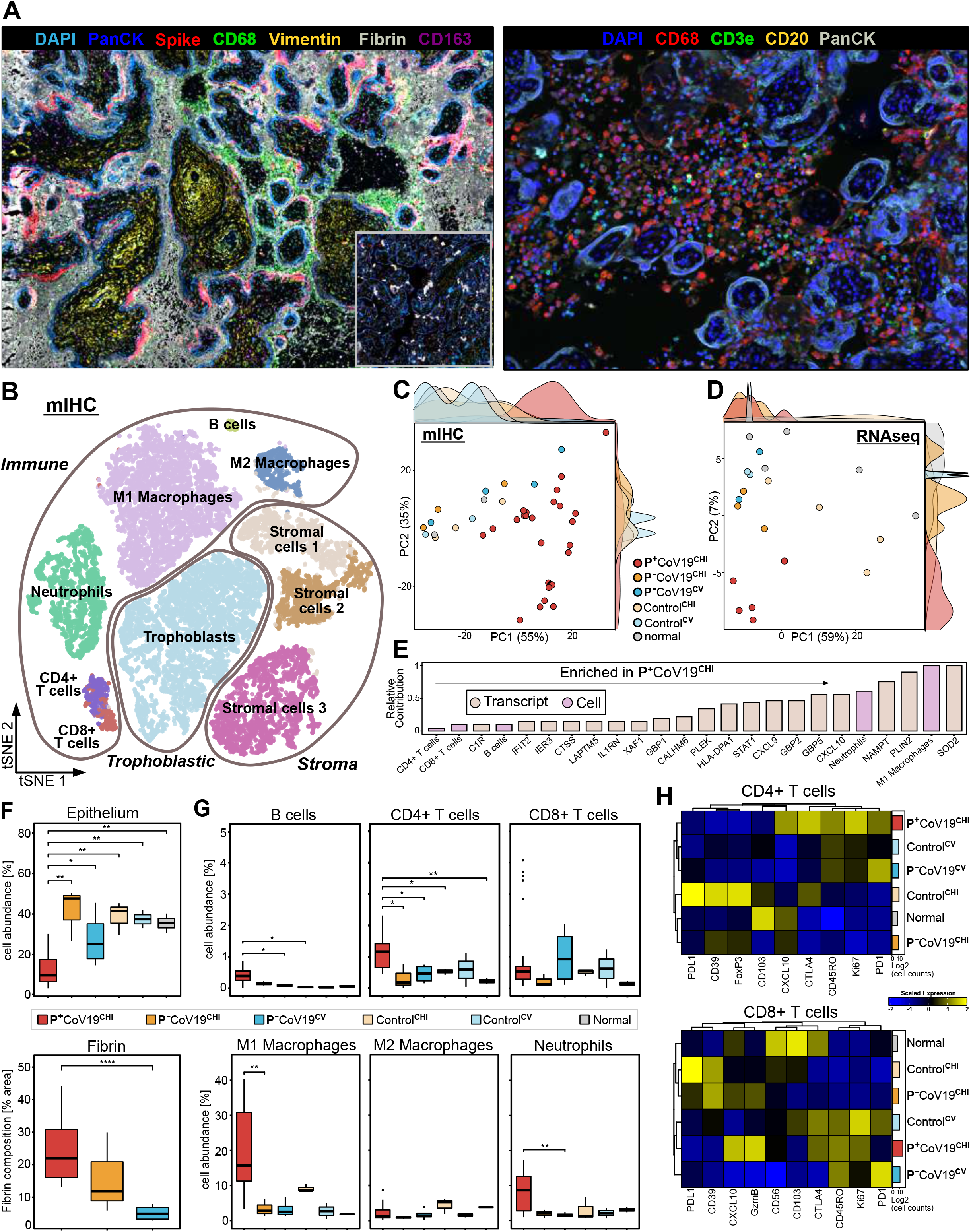
Immune cell landscape of SARS-CoV-2 positive and negative placentas. (A) Representative mIHC images of P^+^CoV19^CHI^ cases highlighting extent of fibrin deposition (left panel) and immune cell composition and location (right panel). (B) tSNE plot showing cell metaclusters identified by quantification of multiplex immunohistochemistry (mIHC) images of placentas from infected mothers and controls. (C and D) Principal Component Analysis (PCA) of cellular abundance of mIHC data (C) and bulk RNAseq transcriptomic reads (D). (E) Combined and normalised PCA loadings of combined mIHC and bulk RNAseq showing relative contribution of immunological and molecular features to P^+^CoV19^CHI^ disease uniqueness. (F) Epithelial cell abundance and fibrin deposition quantification across disease entities. (G) Immune cell abundance in COVID-19 placentitis, control CHI/VUE cases and normal placenta. (H) Functional marker expression on CD4+ T cells (top heatmap) and CD8+ T cells (bottom heatmap) across disease states. See also Figure S3.

For quantitative analysis, mIHC tissue images were divided into smaller regions for deep learningbased segmentation. Three million cells from across all images were segmented with DeepCell then clustered using PhenoGraph ^19^. We initially generated 41 clusters, which were merged into 10 meta clusters (Figure 3B and S3), validated against mIHC images (Figure S3) and visualised on a downsampled tSNE plot (Figure 3B). Principal Component Analysis (PCA) was performed using the abundance of each metacluster to identify features distinguishing different disease states. No batch effects were identified (Figure S3). All regions sampled from the P^+^CoV19^CHI^ cases co-located on the PCA and were distinct from regions sampled from all other disease states and controls suggesting the presence of SARS-CoV-2 is associated with a unique pathological and immunological state; the same pattern was observed when PCA was performed on the bulk RNAseq transcriptomic data (Figure 3C and D).

To identify features contributing to the unique P^+^CoV19^CHI^ disease state, the distinguishing mIHC and RNAseq PCA loadings were calculated, normalised, and combined (Figure 3E). The top discriminating genes included *SOD2* (known to be abundantly expressed by monocytes and neutrophils), and the interferon stimulated genes, *CXCL9, CXCL10, GBP2* and *GBP5. CXCL9* and *CXCL10* are inflammatory cytokines that recruit immune cells, whereas *GBP2* and *GBP5* exhibit broad anti-viral activities and can inhibit furin-mediated processing that is important for SARS-CoV-2 spike maturation^20^. The cell types contributing most to the distinct phenotype of P^+^CoV19^CHI^ cases were M1-like macrophages and neutrophils with a smaller contribution from B-cells and T-cells (Figure 3E).

As expected, based on the earlier qualitative H&E assessment, P^+^CoV19^CHI^ cases also had significantly fewer epithelial cells, consistent with the florid trophoblast necrosis, and elevated fibrin deposition (Figure 3F). Examining the abundance of key immune subsets across all cases, we found that P^+^CoV19^CHI^ had significantly more B-cells and CD4^+^ T-cells with a trend towards increased CD8^+^ T-cells and neutrophils. M1-like macrophages (CD68^+^, CD163^-^) were also significantly more abundant, but M2-like macrophages (CD68^+/-^, CD163^+^) were unchanged (Figure 3G). Assessing the functional status of T-cells in P^+^CoV19^CHI^ showed that CD4+ cells had a distinct phenotype, enriched for CD45RO, Ki-67, CTLA4 and PD1 consistent with proliferating antigen-experienced T-cells. CD8^+^ T-cells in P^+^CoV19^CHI^ were similarly enriched for CD45RO, Ki-67 and CTLA4; however, they lacked PD1 and had high levels of granzyme B consistent with proliferating cytotoxic effector cells (Figure 3H). In normal placentas, CD4+ and CD8+ T-cells were few in number and enriched for CD103, in keeping with a resident memory population (Figure 3H)^21^.

### M1-like macrophages in SARS-CoV-2 positive placentas exhibit increased interactions with T-cells

Given that macrophages have a well-established pathogenic role in SARS-CoV-2 infection^22^ and prompted by our observations that they were not only the most abundant immune cell in virus-positive (P^+^CoV19^CHI^) placentas but that they also co-localised with infected trophoblasts, we examined them in more detail. The COMET mIHC data showed that macrophages in P^+^CoV19^CHI^ placentas were predominantly polarised to an M1-like phenotype (median M1:M2 ratio 14.8, range 0.86-81.2). This was significantly different to normal controls which showed the expected predominance of M2 polarised resident Hoffbauer cells (median M1:M2 ratio 0.48, range 0.47-0.49; Figure 4A). We then examined the compartmental distribution of polarised macrophages, comparing the villous stroma to the intervillous space. Stromal macrophages were almost invariably M2 polarised, consistent with fetal Hoffbauer cells. In contrast, intervillous macrophages in CHI cases, regardless of maternal SARS-CoV-2 status, were predominantly M1 polarised (Figure 4B).

**Figure 4.**
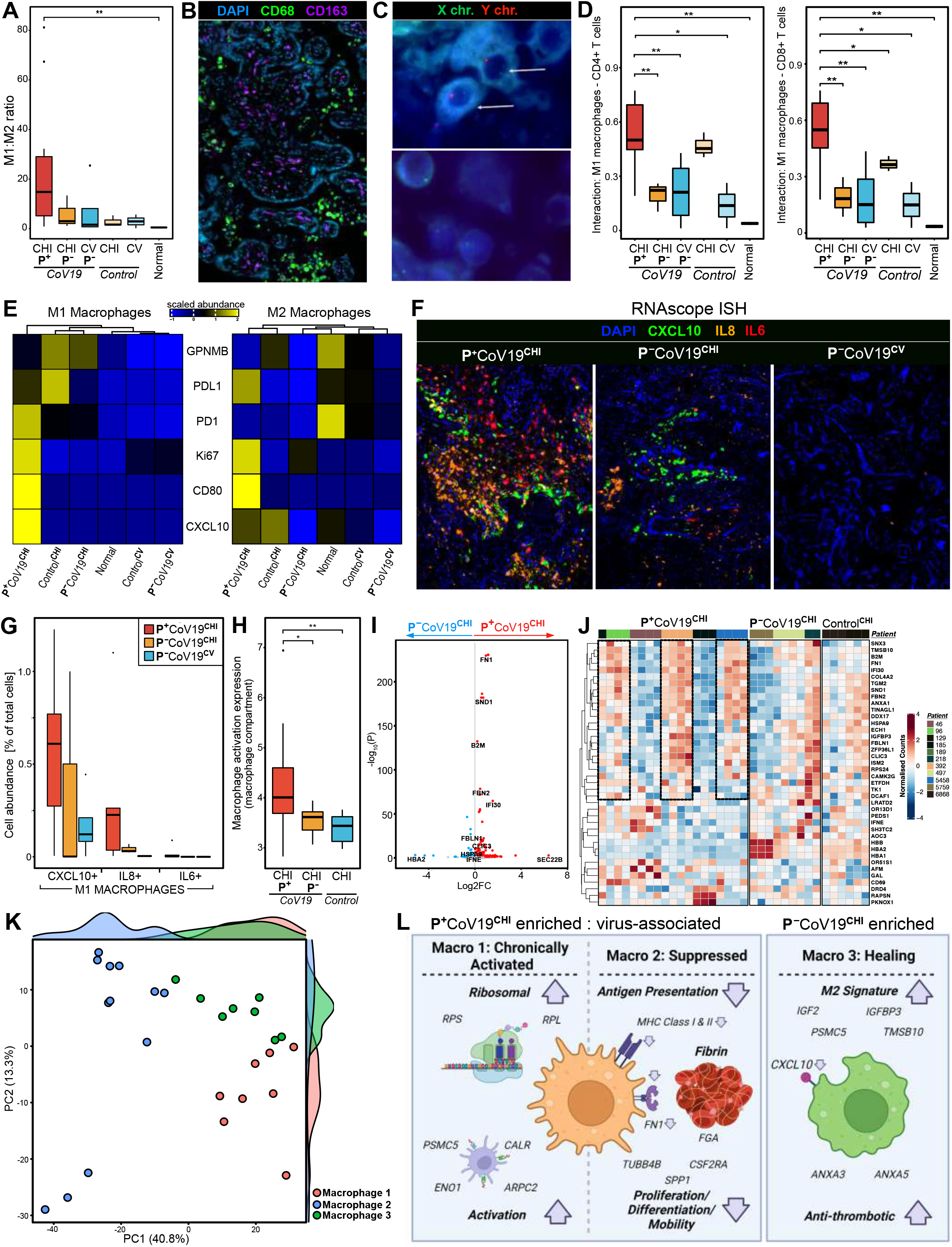
Macrophage characterisation of COVID-19 placentitis. (A and B) Macrophage polarisation ratio of M1 to M2 (A) and localisation of M2 predominately to the villi and M1 predominately to the intervillous space (B). (C) Y chromosome FISH to identify origin of villous and intervillous macrophages (conducted on placentas of male fetuses. Top; villous, Bottom; intervillous). (D) Interaction frequency of M1 Macrophages and CD4+ (left panel)/CD8+ (right panel) T cells. (E) Abundance of M1 macrophages (left panel) and M2 macrophages (right panel) showing positive signal for functional markers. (F and G) RNAscope mISH to identify and localise cytokines *CXCL10, IL6* and *IL8* (F) and quantification of cytokine positive M1 macrophage abundances (G). (H, I and J) Macrophage activation (H) and differential gene expression volcano plot (I) of P^+^CoV19^CHI^ vs. P^-^CoV19^CHI^, followed by expression heatmap of top 30 differentially expressed genes across all disease macrophage compartments (including P^-^CoV19^CV^ and control^CHI^). (K) Unbiased clustering and PCA visualisation of macrophage compartments in P^+/-^CoV19^CHI^ samples. (L) Illustration of three phenotypes of macrophage compartments identified and mechanistic output of each. See also Figure S4.

Given the extensive trophoblast necrosis and the ensuing breakdown in the materno-fetal barrier, we next sought to clarify whether this could have allowed maternal macrophages to infiltrate the fetal compartment or vice versa, contributing to the inflammatory state. To explore macrophage origin, DNA FISH for sex chromosomes was performed for all cases in which the fetus was male allowing maternal/fetal distinction. Across the spectrum of placental diseases, we always observed XY-chromosome signals only in stromal macrophages, confirming their fetal origin, and XX-chromosome signals only in intervillous macrophages, confirming their maternal origin. Therefore, the stromal macrophages were fetal Hofbauer cells, and no cross-compartment mixing was detected (Figure 4C).

To examine macrophage interactions with other immune cell types we generated a Delaunay triangulation mesh from the segmented cell centroids of each mIHC image. Cells that shared an edge in the graph were considered interacting and the likelihood ratio of any two cell types interacting was calculated for each disease state. M1 macrophages in P^+^CoV19^CHI^ more frequently interacted with both CD8+ and CD4+ T-cells than in all other groups including P^-^ Cov19^CHI^ (Figure 4D).

### Macrophages in SARS-CoV-2 positive placentas are the main source of CXCL10 and IL-8

Deeper phenotypic characterisation of macrophages by COMET mIHC demonstrated that M1 macrophages in all placentas with CHI morphology were enriched for the immunoinhibitory proteins GPNMB, PDL1 and PD1 (Figure 4E). However, M1 macrophages in P^+^Cov19^CHI^ cases were unique, expressing CD80, the proliferation marker Ki-67 and the chemokine CXCL10. Ki67, CD80 and PDL1 were also enriched in M2 macrophages from P^+^Cov19^CHI^ placentas, with a slightly increased expression of CXCL10. Thus, both M1-like and M2-like macrophages in P^+^CoV19^CHI^ placentas are phenotypically distinct compared to their counterparts in P^-^Cov19^CHI^ and pre-pandemic diseased and normal controls.

CXCL10 is an important chemoattractant for inflammatory cells, including neutrophils and macrophages, and macrophages are a major source of IL-6 and IL-8. All three cytokines are biomarkers of disease in respiratory SARS-CoV-2 infection^23–25^. Therefore, we performed multiplex RNAscope for CXCL10, IL-6 and IL-8 combined with IHC detection of CD68. SARS-CoV-2 positive placentas (P^+^CoV19^CHI^) expressed high levels of all three cytokines, with lower levels in P^-^CoV19^CHI^ and sparse expression in P^-^CoV19^CV^ (Figure 4F). Quantitative analysis demonstrated that *CXCL10* and *IL-8*, but not *IL-6*, were expressed by macrophages (Figure 4G). Instead, *IL-6* was localised to endothelial cells and spindle-shaped cells, most likely fibroblasts (Figure S4).

### Spatial transcriptomic analysis reveals unique macrophage phenotypes in virus positive cases

Using GeoMx DSP we performed a whole transcriptomic analysis of the macrophage compartment across all COVID-19 disease groups and CHI controls (P^+/-^CoV19^CHI/CV^ and control^CHI^). We first measured macrophage activation, calculating the mean expression of a previously reported macrophage activation gene set (GO:0042116) for each AOI (Figure 4H). Macrophage-rich AOIs showed significantly elevated expression of macrophage activation genes in P^+^CoV19^CHI^ compared with both P^-^CoV19^CHI^ (p=0.014) and control^CHI^ (p=0.009). Focussing on CHI cases from SARS-CoV-2 infected mothers, differential gene expression revealed that, compared with P^-^CoV19^CHI^, P^+^Cov19^CHI^ cases showed significantly increased expression of ECM-associated genes, including *COL4A2*, fibronectin-1 (*FN1*) and fibrillin-2 (*FBN2*), and higher levels of *SND1*, a pro-inflammatory protein that can induce *IL-6* expression (Figure 4I, S4, S5)^26^. Fibronectin-1 was of particular interest given that it complexes with plasma fibrinogen to form fibrin and is known to be expressed by macrophages^27^. Next, we interrogated GeoMx DSP data from multiple spatially distinct macrophage-rich regions from each patient, focussing on the top 20 over- and under-expressed genes. Distinct macrophage phenotypes were evident within P^+^Cov19^CHI^ cases (Figure 4J). To explore these phenotypes in more detail, we unbiasedly clustered the macrophage compartments and visualised them on a PCA plot. Three transcriptionally distinct clusters were identified (Figure 4K). Clusters 1 and 2 were enriched in P^+^CoV19^CHI^ cases. Cluster 1 was characterised by the increased expression of pro-inflammatory genes, including: *RACK1*, which activates the *NLRP3* inflammasome in macrophages^28^; *CCL8/MCP2*, a pro-inflammatory chemokine that is raised in patients with severe respiratory COVID-19 and associated with increased ITU stay; and calreticulin (*CALR*), which drives M1-like macrophage functions and that is required for the development of acute respiratory distress syndrome/acute lung injury (ARDS/ALI) in mouse models^29^. On the other hand, cluster 2 showed a striking down-regulation of genes involved in antigen presentation (*HLA-B, HLA-DRA, HLA-DPB1, HLA-DRB1, B2M, CD74*), M1-polarisation (*CSF2RA, GSTP1*) and fibrin formation (*FBN2, FN1*). Cluster 3 was enriched in P^-^CoV19^CHI^ cases and was characterised by a prominent ‘M2-like’ gene expression signature (*CSF3R, FUS, IFI6*) with over-expression of anti-inflammatory/immunosuppressive genes (*ANXA5, IGF2*) and concomitantly reduced expression of M1-like genes, including CXCL10 (Figure 4L).

### SARS-CoV-2 positive placentas show interferon and anti-viral signatures with localised virus-associated interferon blunting

We next performed a gene set enrichment analysis (GSEA) of the bulk RNAseq data, comparing P^+^Cov19^CHI^ cases to all other disease groups and normal controls (Figure 5A). Gene sets significantly upregulated in infected placentas included *IL-6/JAK-STAT* signalling (consistent with the observed high levels of *IL-6*), allograft rejection, complement activation, coagulation, and hypoxia. The latter two are consistent with the extensive fibrin deposition and mal-perfusion that characterises COVID-19 placentitis. We observed robust interferon-alpha and interferon-gamma signalling in the P^+^Cov19^CHI^ placentas (Figure 5A). Compared to all other groups, placentas from P^+^CoV19^CHI^ cases showed a marked upregulation of interferon-alpha, -beta and -gamma stimulated genes (ISGs, Figure 5B). In particular, 37 *ISGs* experimentally shown to limit SARS-CoV-2 replication^30^ were upregulated in P^+^Cov19^CHI^ (Figure 5C). Moreover, comparing ISGs differentially expressed between virus-positive and -negative cases of COVID-19 placentitis (P^+^CoV19^CHI^ vs P^-^CoV19^CHI^, Figure 5D), we found a striking set of genes implicated in cytokine regulation (*IL1RN*) and antiviral responses (*GBP1, IFIT2*)^31–33^ in the former.

**Figure 5.**
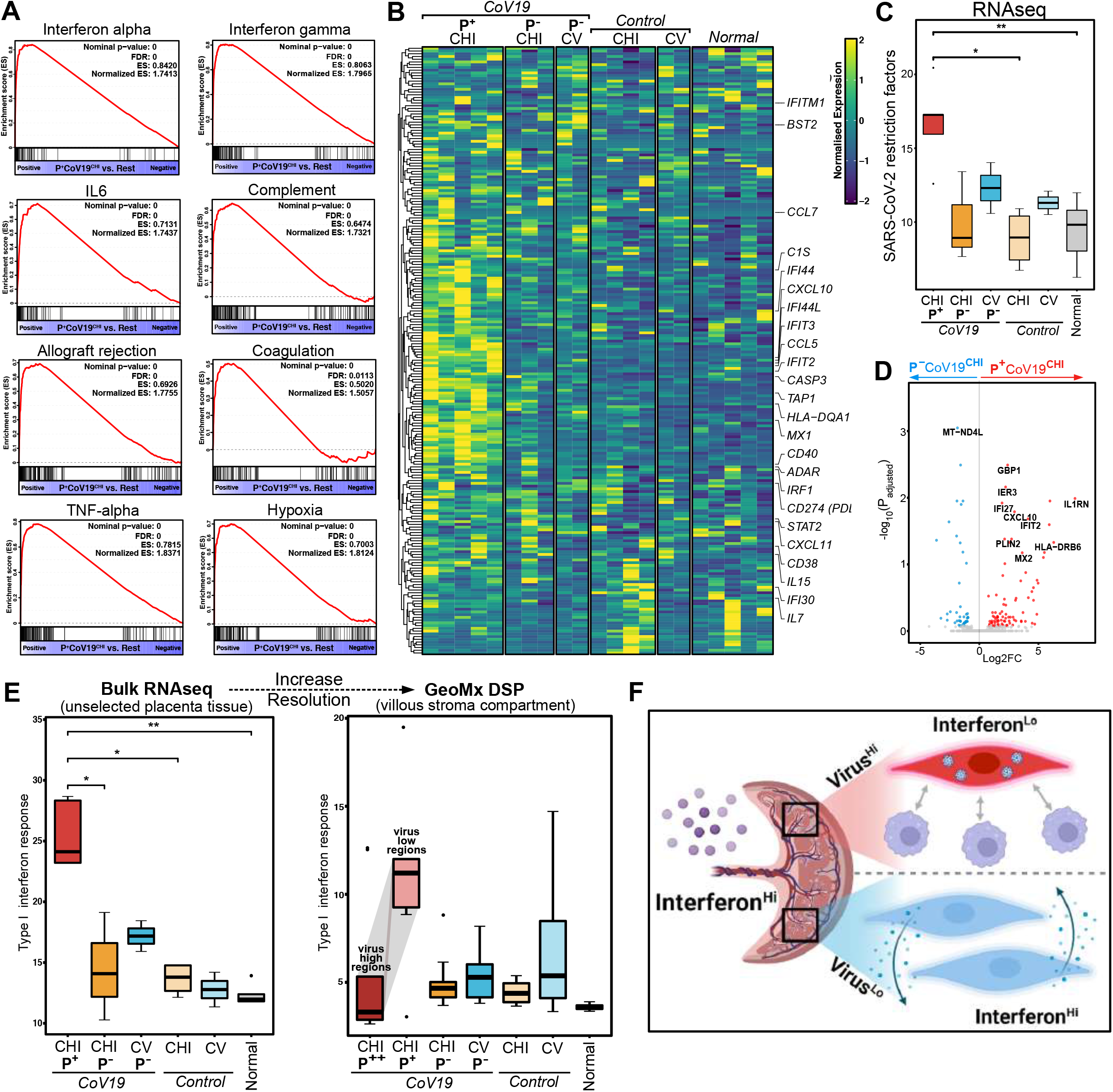
Inflammatory mechanisms within SARS-CoV-2 positive and negative placentas. (A) Gene Set Enrichment Analysis (GSEA) with hallmark gene sets of bulk RNAseq P^+^CoV19^CHI^ vs. rest. (B) Heatmap showing expression of interferon signature genes (type I & II) by bulk RNAseq of placental tissue across all disease states. (C) Bulk RNAseq analysis showing genes differentially expressed between virus positive and negative placentas with CHI pathology from SARS-CoV-2 infected mothers. (D) SARS-CoV-2 viral restriction factor gene set expression by bulk RNAseq across disease states. (E) Interferon Alpha response gene set expression across disease states by bulk RNAseq (left panel) and by villous stroma (VS) NanoString GeoMx DSP compartment with split in P+CoV19CHI cases into VS compartments adjacent to (Virus^Hi^) and not adjacent to (Virus^Lo^) SARS-CoV-2 infected trophoblasts (right panel). (F) Illustration of relationship between virus infection and interferon expression. See also Figure S5.

SARS-CoV-2 is highly sensitive to interferon^34^. To understand how the virus could replicate in placental trophoblasts despite globally high ISG expression we re-examined our GeoMx spatial transcriptomic data. Comparing ISG expression in uninfected stroma and macrophages, adjacent either to infected trophoblast regions or to uninfected trophoblast regions in the same placenta, we found ISGs were markedly down-regulated in the former (Figure 5E, right panel). This contrasted with the tissue-wide elevation of type I interferon response genes seen on bulk sequencing (Figure 5E, left panel). Interferon responses are therefore blunted locally in virus-enriched areas of the placenta despite being upregulated globally.

### The microenvironment of SARS-CoV-2 infected trophoblasts is enriched for innate macrophages and neutrophils but depleted of adaptive B- and T-cells

To explore the VME in more depth we re-analysed our mIHC data, this time calculating how the abundance of each immune cell type varied with increasing distance from SARS-CoV-2 infected trophoblasts (Figure S6). Taking trophoblasts as an anchor point, the abundance of each immune cell type was calculated within concentric circles of increasing radius. To validate this approach, we examined the relationship between infected trophoblast cells and each other. As expected, virus-infected trophoblasts were enriched within the immediate microenvironment of other virus-infected trophoblasts. Compared to non-infected trophoblasts, we found that the immediate neighbourhood of infected trophoblasts was markedly enriched for neutrophils, including a PD1+ subset, PDL1+ M1 and M2 macrophages and CD4+PD1+T-cells (Figure 6A), but depleted of CD8+PD1- and CD4+PD1-T-cells as well as CD20+ B-cells. These findings were confirmed by independent pathology assessment (MP; Figure 6C).

**Figure 6.**
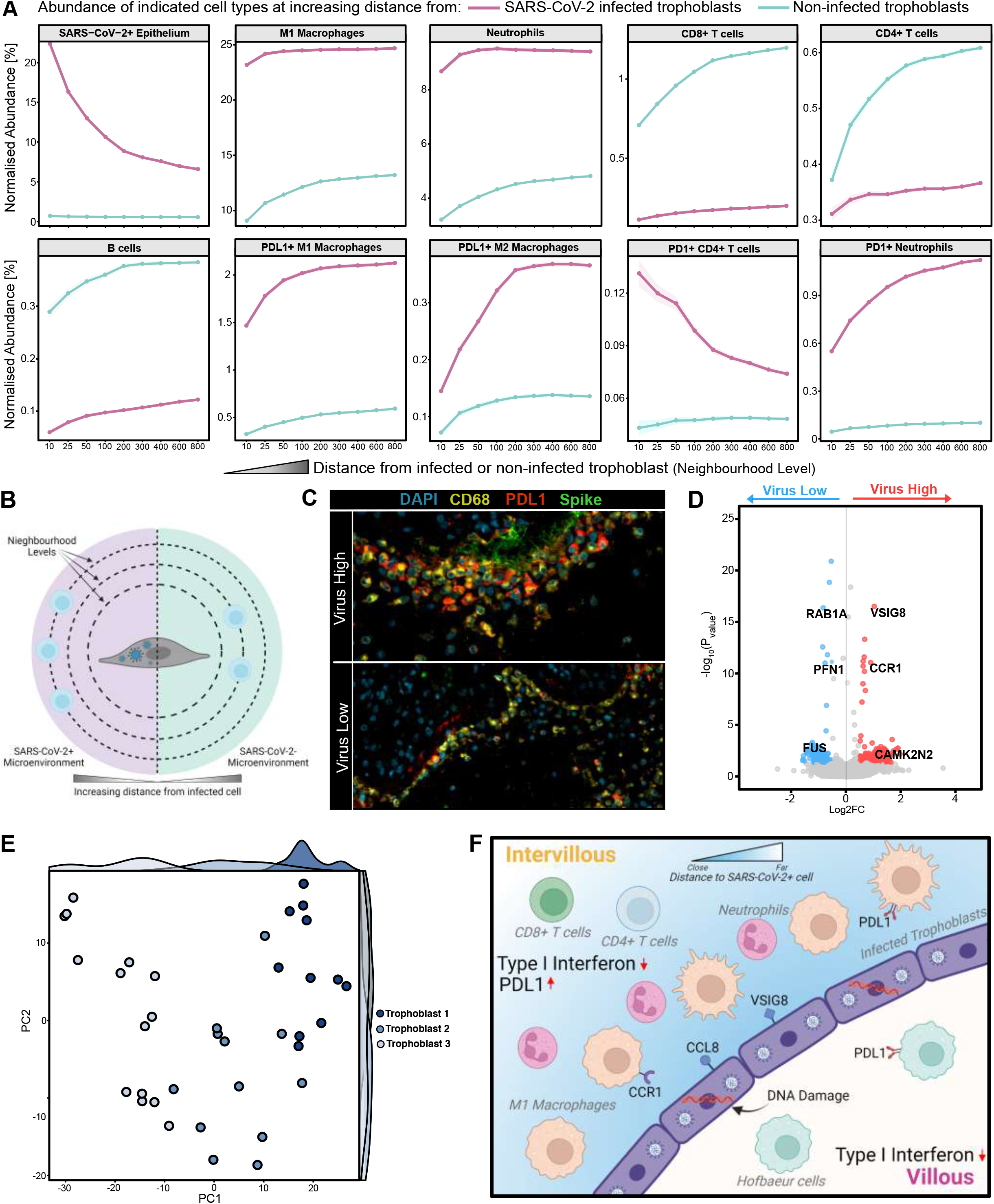
Virus microenvironment of SARS-CoV-2 infected trophoblasts. (A) Normalised abundance of various cell types with increasing distance from SARS-CoV-2 infected trophoblasts. (B) Schematic of virus microenvironment calculation. (C) Representative mIHC images highlighting PDL1 expression in virus high and absence in virus low regions of a SARS-CoV-2 infected placenta. (D) Differential gene expression analysis of SARS-CoV-2 positive vs. negative GeoMx trophoblast compartments. (E) Unbiased clustering and PCA visualisation of trophoblast regions from P+CoV19CHI placentas. (F) Illustration of immune evasion mechanisms operating in SARS-CoV-2 infected placentas. See also Figure S6.

To identify potential drivers of these differences we re-analysed our GeoMx data, first comparing virus-high and virus-low trophoblast ROIs within the same placentas (Figure 6D). Gene ontology of differentially expressed genes showed MHC class I and II antigen presentation pathways were upregulated in virus-high regions. We also observed increased expression of VSIG8, which can bind to VISTA on macrophages to activate an immune checkpoint, and CCR1, a chemotactic factor for macrophages, neutrophils and T-cells that is considered a potential therapeutic target in respiratory COVID-19^35^. Next, we unbiasedly clustered the GeoMx transcriptomic data from trophoblast ROIs of all CHI cases arising in SARS-CoV-2 infected mothers, regardless of virus status (P^+/-^Cov19^CHI^; Figure 6E). This revealed three distinct trophoblast transcriptional signatures. Trophoblast signature (TS)1 was found in virus-positive regions, TS2 in virus-negative regions of both virus-positive and -negative placentas and TS3 in both virus-positive and virus-negative regions. In keeping with a cellular response to infection, TS1 contained stress response genes (*DDIT4*), high expression of *CCL8*, a ligand for *CCR1* and upregulated MHC class I (*HLA-B*). Consistent with the absence of virus, TS2 showed low levels of MHC class I and II genes normally found in trophoblasts. TS3 showed very high expression of *SERPINE1* and *IGFBP3*, markers of epithelial mesenchymal transition (EMT), suggesting a wound healing response^36,37^.

### Multi-omic integration and temporal investigation of COVID-19 placentitis identifies potential viral clearance

To further explore the relatedness of virus-positive and virus-negative cases of COVID-19 placentitis, we integrated the clinical, pathological, and molecular data (Figure 7A). This revealed that P^+^CoV19^CHI^ and P^-^CoV19^CHI^ were clinically and pathologically indistinguishable. However, despite some overlap they could be distinguished at the immunological and molecular levels. In order to resolve the seemingly contradictory overlapping and distinctive features of virus-positive and virus-negative cases of CoV19^CHI^, we considered the possibility that they could represent temporally distinct phases of the same disease. To address this, we first compared their clinical timelines. We found that P^+^CoV19^CHI^ had a shorter clinical course than P^-^CoV19^CHI^, suggesting that P^-^CoV19^CHI^ could represent a more advanced disease stage (Figure 7B). In keeping with this, we found that viral loads and trophoblast numbers also decreased with time (Figure 7C). Collectively, our data indicate that P^-^CoV19^CHI^ cases most likely represent temporal progression of P^+^CoV19^CHI^ associated with virus clearance.

**Figure 7.**
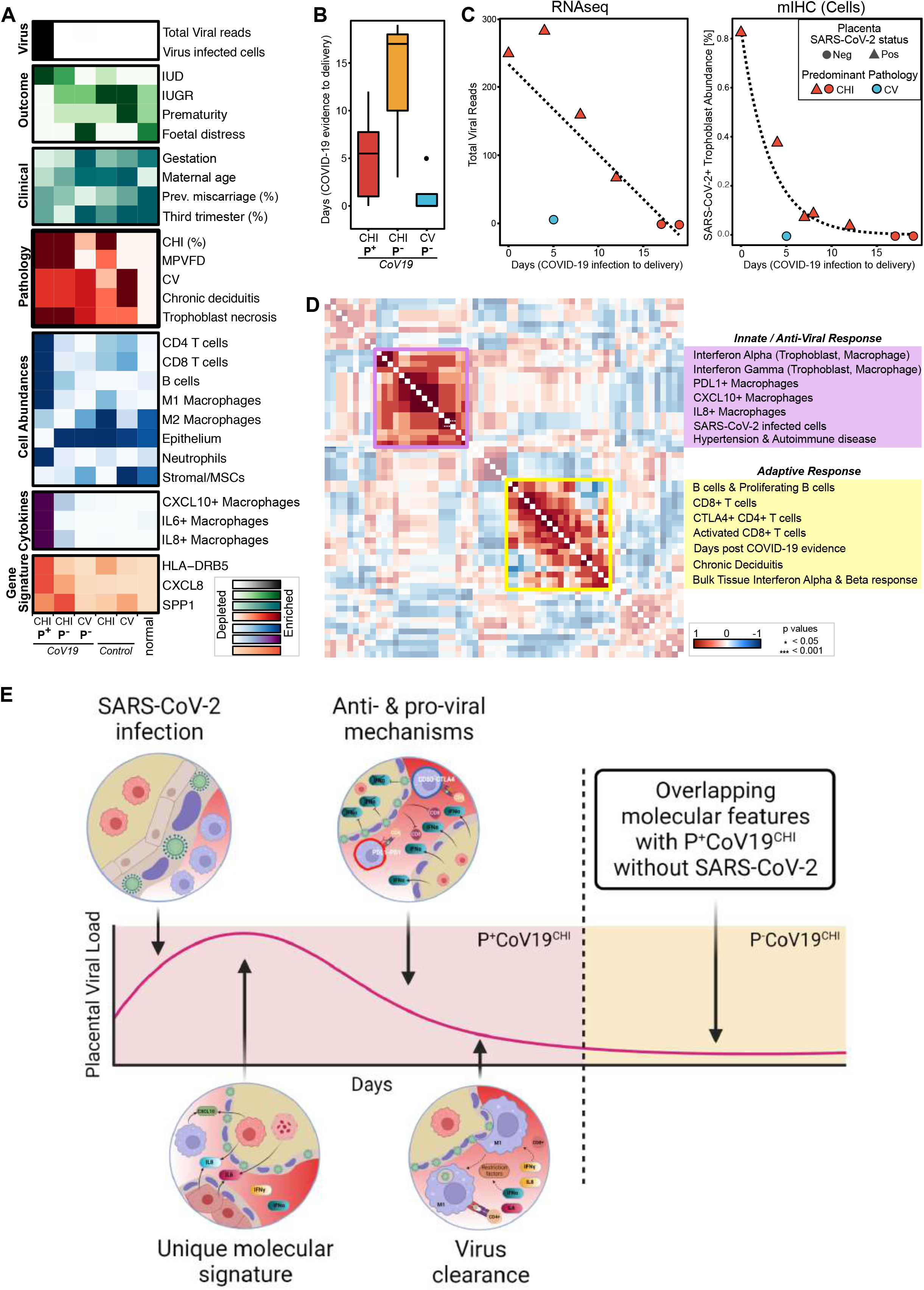
Temporal model integrating the features of COVID-19 placentitis. (A) Summary heatmap of key clinical, pathological, and immunological characteristics of COVID-19 placentitis and control placentitides. (B) Duration of viral infection split by pathological and viral subtype of placenta from SAR-CoV-2 infected mothers. (C) Temporal relationship between placental viral load (left panel) or virus infected cells (right panel) and length of viral infection. (D) Non-parametric correlation matrix of clinical, pathological, and immunological data from P^+^CoV19^CHI^ cases. (E) Temporal model illustration of COVID-19 placentitis. See also Figure S7.

Finally, we generated a non-parametric correlation matrix of the clinicopathological, immunological and cellular features of P^+^CoV19^CHI^ cases (Figure 7D)^38^. We found that innate type responses, including macrophage cytokine expression and compartment-specific interferon-alpha and -gamma responses were consistently correlated. These features correlated with virus burden, which is typically highest early in the disease course. Adaptive-type immune responses, such as immune suppressive CTLA4+ CD4+ T-cells, B cell infiltration and proliferation and CD8+ T cell activation were associated with a more prolonged time course, indicating that these responses develop later in the disease course. Overall, SARS-CoV-2 infection in COVID-19 placentitis represents a unique disease phenotype characterised by an inflammatory transcriptional and cellular signature and several immune evasion mechanisms (Figure 7E & S7).

## Discussion

Whilst COVID-19 placentitis is a known correlate of poor obstetric outcome in the context of maternal SARS-CoV-2 infection, the role direct placental infection plays in mediating the severe pathological changes in the placenta is controversial and poorly understood. Through multi-omic integration of bulk and spatial profiling conducted on placental FFPE tissue from a single cohort of SARS-CoV-2 infected mothers and controls, we reveal multiple inflammatory, anti-viral and immune evasive mechanisms that are contemporaneously activated in SARS-CoV-2 infected placentas. Combined with orthogonal methods of virus detection, we demonstrate a central role for the virus in the pathogenesis of COVID-19 placentitis.

Using a robust multi-omic SARS-CoV-2 detection methodology we localised infection exclusively to syncytiotrophoblasts and showed these cells express both spike and nucleocapsid proteins as well as genomic and sub-genomic RNA, the latter indicating viral replication. Although a subset of maternal (intervillous) macrophages contained viral proteins, these cells were found only in close proximity to infected trophoblasts. Moreover, the localisation of viral proteins in these macrophages was consistent with phagocytosis of virus infected cells rather than direct infection, which has been reported to occur in COVID-19 lungs^39–41^. We did not detect SARS-CoV-2 in any other cell type including stromal or Hofbauer cells reported by others; infection of these cells therefore appears to be infrequent^42^. In keeping with previous reports, we detected placental SARS-CoV-2 infection in only a subset of mothers with respiratory COVID-19 experiencing obstetric complications, all of which exhibited a chronic histiocytic intervillositis (CHI) and massive perivillous fibrin deposition (MFPVD). Similarly, not all placentas from SARS-CoV-2 infected mothers with CHI/MPVFD pathology showed trophoblast infection. However, our data revealed that all CHI/MPVFD cases from COVID-19 positive mothers, including 6 virus positive and 3 negative placentas, showed overlapping molecular signatures. Furthermore, virus loads decreased with increasing disease duration and virus negative cases had longer disease courses compared to virus positive cases. Taken together, the findings suggest a common viral aetiology in both virus positive and negative placentas that is distinct from pre-pandemic CHI. This model would require virus to be cleared from the placenta over time, as has been observed in respiratory COVID-19^17,18^. Collectively, our data suggests SARS-CoV-2 plays a pivotal role in the pathogenesis of COVID-19 placentitis and that direct infection of the placenta is more common than previously appreciated. The findings support the need for a more accurate definition of COVID-19 placentitis based on evidence of clinical infection, pathological features and unique immunological hallmarks which includes placentas without evidence of trophoblast virus infection at the time of pathological assessment. An accurate case definition is essential to monitor disease prevalence in the community and guide public health measures.

Our data also shows a unique immunological signature associated with SARS-CoV-2 positive placentas characterised by counteracting anti-viral responses and pro-viral immune evasive mechanisms, both of which contribute to placental damage. Three key antiviral responses were observed. First, our data shows tissue-wide upregulation of anti-viral type I interferon-stimulated genes (ISG), including known SARS-CoV-2 specific restriction factors^30^. Whilst important for viral restriction, type I interferon responses have also been associated with increased placental damage in animal models for placental Zika infection^5^. Therefore, the anti-viral interferon responses may contribute to placental and fetal demise. Secondly, we found upregulation of classical MHC class-I and class-II expression in trophoblast cells that normally lack these molecules, likely representing a pragmatic but potentially deleterious adaption to aid virus elimination. Combined with transcriptomic evidence of an allograft rejection-type response, the findings are indicative of a potential breakdown of maternal immune tolerance to the placenta. An allograft rejection-like response due to loss of immune tolerance has been postulated to play a role in non-virus associated CHI due to similarities with the immune responses in transplant-associated rejection^43^. Thirdly, we found an enriched infiltration of phagocytic M1 macrophages palisading around infected trophoblasts that secrete soluble fibrinogen, the precursor to insoluble fibrin. These cells likely contribute to the extensive fibrin deposition that is a hallmark of COVID-19 placentitis and compromises placental perfusion and viability. In turn fibrin, along with interferon-γ, has previously been shown to upregulate the neutrophil and macrophage chemoattractant *CXCL10* in macrophages, potentially forming a positive feedback loop^44^. Macrophages were the dominant source of CXCL10 in virus positive placentas. Intriguingly, *CXCL10* has been associated with reduced immune clearance of alphavirus disease in animal models and represents a potentially druggable therapeutic target^45^. *CXCL10* release from macrophages is inhibited by statins, a widely available drug which can be used in the later stages of pregnancy^46^. This strategy could target the fibrin-mediated marcrophage-*CXCL10* positive feedback loop and enhance viral clearance if cases could be identified at an early stage. *CXCL10* expressing macrophages are also abundant in the lungs of patients with SARS-CoV-2 respiratory infection and *CXCL10* has been proffered as a therapeutic target in this context^44,47^.

Pro-viral immune evasive mechanisms were localised to the immediate virus microenvironment and were only detectable due to our high-dimensional spatial profiling approach. They included the establishment of an immunosuppressive niche, enriched for PDL1 expressing M1 macrophages and PD1+ CD4+ T-cells, and depletion of and CD8+ cytotoxic T-cells, adjacent to SARS-CoV-2 infected trophoblasts. Levels of ISG expression were also decreased in these regions, potentially explaining how the virus can replicate in the placenta despite globally high ISG expression. The mechanism responsible for localised interferon blunting is currently unclear. Although type III interferons are constitutively expressed by the syncytium, ISG induction in villous explants occurs in response to type I interferons but not type III interferons^5^. Therefore, the localised reduction of ISGs we observed may reflect reduced type I interferon levels in the vicinity of infected trophoblasts. The source of type I interferons in SARS-CoV-2 infected placentas, their potential contribution to placental inflammation and harm, and how interferon responses are blunted in virus positive regions are priorities for future work.

It remains unclear why the virus infects the placenta in only a minority of mothers who contract respiratory COVID-19 infection. Overall, the proportion of mothers infected with SARS-CoV-2 who show placental or fetal involvement is low (<2%); however, the most severe obstetric complications are concentrated amongst those mothers with COVID-19 placentitis^48^. Notably, there is no correlation between respiratory disease severity and obstetric outcome. Most of the stillbirths observed in our study, and the wider literature, were associated with asymptomatic or mild COVID-19. Several studies, including our own, show that trophoblasts consistently express ACE2, the receptor for SARS-CoV-2, along with other proteins involved in viral entry^49^. We and others have also shown that the distribution of ACE2 in un-infected placentas, including prepandemic CHI, CV and normal controls, is restricted to the stromal side of the trophoblast layer and therefore presumably inaccessible to virus via the maternal circulation^50,51^. In contrast, our data revealed that SARS-CoV-2+ placentas, as well as SARS-CoV-2-placentas with features suggestive of prior viral infection, showed ACE2 expression localised to the maternal circulatory side of the trophoblast layer. ACE2 localisation to maternal circulatory side of the trophoblast layer may explain why some women are more susceptible to direct placental infection by SARS-CoV-2.

CHI/MPVFD occurring before the pandemic, and therefore not associated with SARS-CoV-2, occurs at low incidence in pregnancy but has a 25-100% chance of recurrence in subsequent pregnancies^43^. None of the CoV19^CHI^ mothers in our cohort had prior history of CHI. This observation further strengthens the case that the virus plays a key role in COVID-19 placentitis rather than opportunistically infecting placentas that were developing CHI. Our data showing evidence of loss of maternal tolerance to placenta in COVID-19 placentitis, as previously observed for non-virus associated pre-pandemic CHI^43^, is important. It provides a mechanistic basis for recurrence in the absence of infection in subsequent pregnancies. Close obstetric followup, monitoring and counselling should therefore be offered to patients and studies to record the true risk of recurrence should be implemented. Our findings also emphasise the need for ongoing public health measures to safeguard vulnerable pregnant women from the complications of placental SARS-CoV-2 infection, given the pivotal role of the virus in driving the unique immune landscape of COVID-19 placentitis.

## Acknowledgements

This work was funded by the Science Foundation Ireland (SFI) COVID-19 Rapid Response Grant (DISECT project; 20/COV/8544) and the Medical Research Council (MR/T001755/1). HJH and ZS are funded by a Medical Research Foundation intermediate career fellowship to ZS (UKRI, MRF-169-0001-F-STAM-C0826). We acknowledge the technical support of the University of Birmingham Genomics Facility and Birmingham Tissue Analytics facilities expertise in spatial profiling. We thank Lunaphore Technologies for the provision of a COMET multiplex immunohistochemistry system and associated reagents. All illustrations were created with BioRender.com

## Author Contributions

Conceptualization: M.P, E.F, P.G.M, G.S.T.; Methodology: M.P, C.I.L., E.F; Software: E.F.; Validation: M.P, C.I.L., T.P., G.R.; Formal Analysis: E.F., M.P.; Investigation: M.P, C.I.L., T.P., G.H., G.R.; Resources: K.J.H, B.E.W., Z.S., H.J.H., B.H., T.M.; Data Curation: M.P., E.F., B.H., T.M.; Writing - Original Draft, M.P, E.F, C.I.L, P.G.M, G.S.T; Writing - Review & Editing, M.P, E.F, B.H., T.M., C.A.T., P.G.M, G.S.T.; Visualisation: M.P., E.F., G.S.T.; Supervision: N.S., S.D.D., P.G.M., G.S.T.; Project Administration: M.P.; Funding Acquisition: M.P., E.F., P.G.M & G.S.T;

## STAR Methods

### RESOURCE AVAILABILITY

#### Lead Contact

Information and requests for resources and/or reagents are to be directed to the Lead Contact, Dr. Graham S. Taylor.

#### Materials Availability

This study did not create any unique reagents.

#### Data and Code Availability

Data and code will be made available upon publication.

### STUDY COHORT & SAMPLING

#### Patient selection, tissue sampling and processing

Placental formalin fixation and paraffin embedded (FFPE) tissue samples from SARS-CoV-2 positive mothers suffering obstetric complications in the West Midlands, UK, were referred to the perinatal pathology department at Birmingham Women’s and Children’s Hospital for pathological assessment (n=13). Macroscopic and morphological assessment was performed by paediatric and perinatal pathologists B.H. and T.M at the Cellular Pathology Department of Birmingham Women’s and Children’s Hospital. Tissue was selected and sampled based on macroscopic appearance for formalin fixation and paraffin embedding and subsequent morphological assessment on haematoxylin and eosin (H&E) stained sections along with CD68 immunohistochemistry. Anonymised FFPE tissue was transferred to the University of Birmingham for further laboratory experiments under the LoST-SoCC study (IRAS 193937) following approval by the research ethics committee (19/NE/0336). Diseased control placentas of mothers suffering obstetric complications pre-pandemic were also sampled, totalling 24 tissues. Maternal respiratory infection was confirmed in 12/13 cases by SARS-CoV-2 nasopharyngeal swab PCR; the mother of the remaining placenta returned a negative swab but was COVID-19 symptomatic and showed SARS-CoV-2 infection of the placenta. Upon pathological assessment (B.H. & T.M.) of the control tissue, 4 cases were classified pathologically as CHI, 2 as VUE and 5 as normal. Curls and slides were constructed from the tissue blocks. For COMET mIHC imaging, 4.5 mm tissue sections were extracted from blocks and embedded into an arrangement which covered a 9 × 9 mm window (2 × 2 tissues per slide). Slides were subsequently cut onto positively charged frost-free slides in preparation for chromogenic and mIHC staining. Serial sections of each slide were stained with H&E and scanned with an Aperio CS2. Clinical features were extracted from the clinical data of all mothers, while pathological features were annotated retrospectively by pathologists (M.P, TM, BH; SI Table 1). Full details of the patient and clinical information can be found in SI Table 1.

### METHOD DETAILS

#### Chromogenic Immunohistochemistry

FFPE sections, 5μm thick, were baked for 3 hours prior to staining to ensure tissue adherence. The Leica BondMax IHC protocol F (DAB) was used with the following conditions: bake and dewax, epitope retrieval 2 for 20 minutes and primary antibody incubation for 1 hour. Slides were subsequently mounted with DPX mountant media and scanned (Aperio CS2).

#### COMET Multiplex Immunofluorescence Biomarker Panel & Staining

A 29-plex biomarker panel was designed and validated to identify cell types present in the placenta as well as quantify functional states and signalling molecules associated with COVID-19 placentitis. Slides were de-waxed, baked and antigen retrieved (102°C for 1 hour in a pH 9 buffer) using the PT module (Thermofisher) before a brief wash in multi-staining buffer. The slides were then loaded into the COMET stainer (Lunaphore Technologies). Before antibody staining cycles commenced, blank images of the TRITC and Cy5 channels were captured for autofluorescence subtraction during image processing post acquisition. Primary antibodies were diluted to the desired concentrations (see SI Table 2) in multi-staining buffer and incubated for 4 minutes in each cycle. Secondary antibody incubation was carried out for 2 minutes with a 2-minute elution. Exposure times used were 50ms for DAPI, 400ms for the TRITC channel and 200ms for Cy5. Secondary antibodies in the TRITC channel (mouse antibody specific) were diluted to a concentration of 1:100 while Cy5 channel secondaries (rabbit antibody specific) were diluted to 1:400. DAPI images were taken in every staining cycle to ensure alignment of cycles during image processing. All images were acquired with a 20x objective. All antibodies were optimised using negative staining (secondary antibody only) and elution/re-staining step protocols.

#### RNAscope in situ hybridisation/immunohistochemistry

Two panels utilising RNAscope ISH protocols were implemented to detect sub-genomic RNA and cytokine producing cells, respectively. In the first panel, SARS-CoV-2 and nCoV-2019-S Sense probes were used in a 2-plex panel to detect the virus and replicating virus respectively. In the second panel, a 3-plex protocol (CXCL10, IL8, IL6) was used to detect cytokine expression across the cohort. Both panels were combined with immunohistochemical detection of CD68. The RNAscope ISH protocols were conducted as specified by ACD (Cat. No. 320293). Briefly, FFPE slides were baked at 60 °C for 1 hour before being deparaffinised in xylene and dehydrated 100% ethanol. Slides were then dried before H2O2 was added for 10 min at room temperature (RT). Target retrieval was achieved by incubating the slides in boiling antigen retrieval buffer (<98°C) for 15 minutes. Slides were then treated with Protease Plus for 15 minutes at 40 °C. For panel 2, C2 and C3 probes were diluted in C1 probes (C2 probe was diluted in C1 for panel 1) at a ratio of 1:50 and were added to the slides for 2 hours at 40 °C. C1 probes were detected with Opal-520 (FP1487001KT, Akoya Biosciences), C2 probes with Opal 620 (FP1495001KT, Akoya Biosciences) and C3 probes with Opal 690 (FP1497001KT, Akoya Biosciences). For IHC detection of CD68, post-RNAscope slides were boiled in citrate buffer for 15 minutes, blocked with casein for 10 minutes and incubated with anti-CD68 primary antibody (clone KP-1, Dako) for 1 hour at RT. Primary antibody was detected using universal HRP secondary antibody (MP750050, Vector Labs) and Opal 780 (FP1501001KT, Akoya Biosciences). Before mounting with Prolong Gold, the slides were counterstained with DAPI. Slides were imaged on a Vectra Polaris automated imaging system.

#### Fluorescent in-situ hybridisation for X/Y chromosomes

Unstained placental sections on positively charged slides were baked at 60°C overnight prior to deparaffination with fresh UltraClear, then cleared using 100% methanol and distilled water. The sections were immersed in 0.2M HCl for 23 minutes, washed using distilled water and placed in the heat pre-treatment solution from Cytocell’s LPS100 Tissue PreTreatment Kit for 50 minutes at 95°C. The slides were washed with distilled water and a few drops of the enzyme digestion solution was added to the sections and incubated at 37°C for 40 minutes. The Cytocell LPE0XG and LPE0YqR FISH probes and sections were denatured and hybridised together according to the manufacturer’s guidelines. The tissue was counterstained with DAPI obtained from Cytocell and analysed using the Olympus BX51 fluorescent microscope with images captured using Metasystems’ Isis software.

#### NanoString GeoMx RNA Spatial Transcriptomics

FFPE TMAs specifically created for GeoMx DSP analysis (16 tissue samples on 4 slides comprising 5 P^+^CoV19^CHI^, 3 P^-^CoV19^CHI^, 2 P^-^CoV19^CV^, 2 control^CHI^, 2 control^CV^ and 2 normal placentas) were stained with pan-cytokeratin, CD68, SARS-CoV-2 nucleocapsid protein and DAPI to identify trophoblasts, macrophages and SARS-CoV-2 infected cells. Regions Of Interest (ROI) were subsequently selected based on this staining by a registered pathologist (M.P.) and assigned as to one of four compartments: villous stroma, decidua, trophoblast or macrophage. Virus positivity of each region was also assigned by a registered pathologist (M.P.). The NanoString Whole Transcriptome Atlas (WTA) gene panel was used^52^. Oligonucleotides were cleaved off the antibodies and gathered within each ROI as per the GeoMx DSP protocol. The photocleaved oligos were sequenced on an Illumina NextSeq 550.

#### QuantSeq Library Prep & Sequencing

RNA extraction was carried out using the RNeasy FFPE Kit (Qiagen 73504) according to the manufacturer’s instructions for samples containing more than 2 curls. Briefly, curls were deparaffinized in xylene and rehydrated with ethanol washes. The tissue was digested in Buffer PKD with 10 μL of proteinase K and 2 μL of β-mercaptoethanol at 56°C for 15 min followed by incubation at 80 °C for 15 min. Samples were cooled on ice for 3 min and the digested tissue was pelleted by centrifugation. DNA was digested in DNase Booster Buffer with 10 μL of DNase I for 15 min at room temperature. The binding conditions of the sample were adjusted by adding buffer RBC and ethanol. The RNA was applied to a RNeasy spin column and washed with buffer RPE before eluting in 30 μL of RNase-free water. 5 μL of RNA was removed to determine the quality and quantity of the extracted RNA using the Agilent 4200 TapeStation and Qubit, respectively. The remaining 25 μL was utilised for QuantSeq library prep. Library prep was completed using the Lexogen QuantSeq 3’ mRNA-Seq Kit FWD for Illumina. QC was monitored by spiking in 0.5uL of SIRV to allow technical evaluation of library prep and sequencing performance. The prepared RNA was sequenced on an Illumina NextSeq 550.

#### SARS-CoV-2 inection of VERO cells

SARS-CoV-2 hCOV-19/England/2/2020 was provided by Christine Bruce, UK Health Security Agency, Porton Down, UK and passaged once in Vero-TMPRSS2 cells, provided by Jane McKeating, University of Oxford. Vero cells were infected for 48 hours at MOI 0.04 and fixed with neutral buffered formalin. Cells were then pelleted by centrifugation at 600rpm for 10 minutes and processed through alcohol and xylene to paraffin wax and embedded. Infection was detected with rabbit anti-spike antibody clone CR3022 (The Native Antigen Company, Oxford UK) or mouse anti-nucleocapsid antibody clone CR3018 (Absolute Antibody Ltd, Cleveland UK) and cell nuclei were visualised with Hoechst 33342 (Thermo Fisher Scientific). Infection was imaged using the Cellinsight CX5 HCS Platform (Thermo Fisher Scientific).

#### Virus Variant Identification

Analysis of SARS-CoV-2 spike mutations was performed by RT-PCR. RNA, purified from FFPE scrolls using a Qiagen RNeasy FFPE kit, was amplified using Qiagen OneStep RT-PCR reagents and SpikeSNP Ext and R1 primers^53^. Cycling conditions were 95C 15’; 40 cycles of 94C 20s, 58C 20s, 72C 40s; and then 72C 7’. Amplicons 350bp in size were purified from agarose gels using a Qiagen gel extraction kit and Sanger sequenced by Source Bioscience (Nottingham UK) using the SpikeSNP Ext and R1 primers.

### QUANTIFICATION AND STATISTICAL ANALYSIS

#### COMET mIHC Image Processing and Cellular Segmentation

After acquisition, images were shading corrected, cycle aligned, stitched and merged into OME-TIFF pyramidal format within the Lunaphore COMET Explorer software. Autofluorescence subtraction for TRITC and Cy5 was conducted in the Lunaphore COMET Viewer v2. Single channels from each OME-TIFF stack were then extracted and TMAs were de-arrayed and saved as individual channel tiff files. The full-face slide sections imaged were split into quadrants in order to run through a deep learning segmentation model. Lineage marker channels (CD45, pan-cytokeratin, vimentin, aSMA and CD15) were merged into a ‘pseudo-membrane’ image and tiled to 256 × 256 pixel images. These tiles were used as input to the Mesmer model of the DeepCell segmentation library^54^. Default settings for the Mesmer model were used and inference was made on both the membrane and nucleus channels. After inference, labelled cells were contracted by one pixel to ensure no cells were touching and subsequently interpolation-based stitching was used to build back up to the original image size. The image was binarised to negate the effect of interpolation on labelled regions and then re-labelled using the ‘label’ function of scikit-image. Labelled cells were then expanded back up by one pixel to return to their original size. To extract nuclear information from nuclear channels (DAPI, FoxP3, MUM1 and Ki67), each membrane label was connected with the most appropriate nucleus label using the ‘find_nuclear_label_id’ function from ARK^54^. The ‘region_props’ function from scikit-image was then used to extract mean channel intensities as well as other morphological features and saved in an FCS format. Labelled mask images were saved as tiff files.

#### Clustering and Cell Phenotype Identification

FCS files generated from COMET imaging were imported into MISSILe. Expression values were compressed to the 99.9th percentile per region. The morphology metrics cell size, solidity and eccentricity were z-scored and used as initial quality control steps (−2 < cell size < 2, solidity > −2, eccentricity < 1.95). Clustering was performed with Phenograph through the FastPG package in R^19^. Two separate clustering runs were conducted: low resolution and a subsequent re-clustering. Only lineage markers were utilised for low resolution clustering to identify main cell phenotypes. Dot plots for the enrichment and expression of each marker per cluster were constructed and clusters were merged to form meta clusters (see Figure S3; conducted by experienced single cell researchers E.F. & G.S.T.). Clusters were validated by plotting the centroid of each phenotype over the original images and qualitatively assessing matches (Figure S3). To assess batch effects between slides/regions, PCA was run on the phenotype percentages per region. The results were coloured by slide and disease state and qualitatively assessed as to the influence of each on the cell type abundance (Figure S3). Each meta cluster was re-clustered with all functional and immune checkpoint markers and subsequently annotated. Abundances of each cluster were calculated as a percentage of total detected cells as well as per unit of tissue area. For qualitative assessment of the spatial orientation of cell phenotypes, the centroids of each cell were plotted by region of interest and coloured according to cluster annotation (Figure S3).

#### SARS-CoV-2 Infected Cell Identification

To ensure correct identification of SARS-CoV-2 infected cells, only cells positive for both SARS-CoV-2 spike and nucleocapsid antibodies by clustering were denoted as infected. Cases with positive cells by COMET mIHC were checked for expression of RNAscope SARS-CoV-2 probes and viral reads to the SARS-CoV-2 transcriptome by bulk RNAseq to ensure correct identification of virus infected cases (Full validation in Figure 2 & S2).

#### Cellular Interactions and Neighbourhoods

To quantify the interaction of cell phenotypes, a Delaunay triangulation graph was calculated using the centroids of each cell. Vertices that were connected as well as phenotype of that vertex were recorded. The likelihood ratios and relative frequencies of all interactions were subsequently calculated and visualised with boxplots split per disease state^55^. Cellular neighbourhoods were calculated by recording the 10 nearest neighbours of each cell and clustering these using k-Nearest Neighbours (kNN) with k = 10 as previously described^55^. Distances between two cell types of interest were calculated with the nn2 function of the RANN library.

#### Virus microenvironment

To quantify the immediate immune microenvironment of SARS-CoV-2 infected trophoblasts, cell phenotype abundances of each phenotype were recorded as a function of distance from the infected cells ^56^. The nn2 function of the RANN library was used to calculate the N (where N = 10, 20, 40, 80, 100, 200, 400, & 800) number of cells closest to a SARS-CoV-2 infected trophoblast. The phenotype of each cell was then calculated as a percentage of the total cells in each bin (N) and plotted. For fair comparisons, the microenvironments of virus infected trophoblasts were compared to the number of cells matched microenvironments of non-infected trophoblasts within the same samples.

#### RNAscope ISH Image Processing and Analysis

After RNAscope ISH slide acquisition, images were tiled in Phenochart and spectrally unmixed in inForm. After unmixing, tiles were stitched and subsequently segmented in QuPath^57^. The intensity of each channel in each cell was extracted using the ‘Measurements’ functionality. These intensity measurements were concatenated and imported into R. Channels were thresholded by a pathologist (M.P.) and macrophages (denoted by positive CD68 threshold) were assessed for percentage of positive expression of each cytokine (*CXCL10, IL6, IL8*). The percentage of cytokine positive macrophages were calculated as a function of total cells and total macrophages and were plotted.

#### Processing and Analysis of GeoMx DSP RNA Spatial Profiling

Fastq files from the GeoMx DSP sequencing run were concatenated and processed through the NanoString GeoMx NGS Pipeline to generate Digital Count Conversion (DCC) files. DCC files were imported into the NanoString interactive data analysis and visualisation software. 3 regions were excluded from the analysis through the segment and probe QC steps and Q3 normalisation was subsequently performed. Gene count and metadata files were exported in csv format and imported with annotated metadata files into R. Gene set expression was calculated by taking the addition of genes within selected gene sets per region. Differential expression between disease states within each compartment was conducted with glmmSeq^58^, using age and gestation as fixed effects and sample ID as random effect.

#### Bulk RNAseq Alignment and Analysis

Fastq files generated from each flow cell were concatenated for each sample replicate. RNA quality control of the fastq files was then conducted with FASTQC ^59^. Subsequently, according to Lexogen recommendations of the 3’ assay, the adapter contamination, polyA read through and low-quality tails were trimmed using bbduk. The SARS-CoV-2 genome files from NCBI were merged with the GRCh38 Homo sapiens genome, and a new reference was created with STAR aligner^60^. Alignment was then conducted using STAR, with duplicate reads subsequently removed with PICARD. The resulting BAM files were indexed using SAMtools and counted using HTSEQ-count ^61,62^. The count files were then imported into R and subsequently DESeq2, where they were normalised using the median of ratios ^63^. Differential expression was conducted within DESeq2 with maternal age and gestation as covariates. Gene set enrichment analysis was conducted with the GSEA program from the Broad Institute ^64,65^, heatmaps were visualised using the ComplexHeatmap library and differentially expressed genes were visualised using volcano plots from EnhancedVolcano ^66^. Gene set expression was calculated by taking the addition of genes within selected gene sets per region.

#### Dimensionality Reduction Analysis

Principal Component Analysis (PCA) was conducted using the prcomp function in R. For PCAs of gene expression, the top 500 most variable genes were used as input. Multiple Correspondence Analysis (for pathological and clinical variables) was calculated with the MCA function of FactoMineR in R. Both PCA and MCA were plotted using ggplot2.

#### Correlation matrices

The non-parametric correlation matrix was calculated as previously described and plotted with corrplot (Figure 6D)^38^.

#### Variant/Cases Epidemiological Analysis

Data for the epidemiological analysis was downloaded from worlddata.info for specific regions of the UK relevant to this study. Case numbers, dominant variants, and vaccine uptake (first and second dose) were plotted as a function of the date.

## Supplementary Figure Legends

**Figure S1. Detailed experimental, clinical, and morpho-pathological features, related to Figure 1**

(A) Summary of numbers of placenta samples analysed by the different multiomic platforms.

(B) Multiplex immunohistochemistry (mIHC) antibody panel used for COMET mIHC analysis of FFPE tissues.

(C) Region of Interest selection strategy to analyse different placental tissue compartments on the GeoMx Digital Spatial Profiler (DSP). The NanoString Whole Transcriptome Analysis (WTA) panel was used to measure mRNA expression.

(D) SARS-CoV-2 positive cases sampled plotted over total COVID-19 cases in the Birmingham area.

(E) Multiple correspondence analysis of placental pathological features annotated by pathologists M.P., B.H. & T.M.).

(F) Macroscopic morphology of SARS-CoV-2 positive placentas showing diffuse lesional involvement affecting the whole placenta

(G) Haematoxylin and eosin-stained images of COVID-19 associated CHI showing common morphological features.

(H) CD68 chromogenic immunohistochemistry of COVID-19 (top panel) and control CHI (lower panel).

**Figure S2. Validation of SARS-CoV-2 detection and viral variant analysis, related to Figure 2**

(A) Validation of SARS-CoV-2 spike and nucleocapsid antibodies.

(B) Validation of SARS-CoV-2 spike sense and anti-sense RNAscope fluorescent ISH probes.

(C) Qualitative validation of virus infected cell identification from mIHC images.

(D) SARS-CoV-2 positive mothers placental sample date plotted over dominant viral strain in the UK.

(E) PCR and Sanger sequencing of the spike gene from two P^+^CoV19^CHI^ placental samples.

(F) Known and potential SARS-CoV-2 entry gene expression of NanoString GeoMx DSP trophoblast compartments.

**Figure S3. Clustering, validation, and neighbourhood analysis of COVID-19 placentitis, related to Figure 3**

(A) Dot plot of metacluster antibody expression by multiplex immunohistochemistry (mIHC).

(B) Examples of metacluster validation projected over mIHC images.

(C) Spatial point pattern of P+CoV19CHI vs. ControlCHI showing the tissue architecture damage seen on H&E.

(D) Principal Component Analysis of cellular abundances coloured by slide of origin.

(E) Tissue neighbourhood enrichment heatmap.

(F) Visualisation of tissue neighbourhoods for P+CoV19CHI, P-CoV19CHI and. ControlCHI

(G) Abundance (%) of each neighbourhood across each disease state.

**Figure S4. Cytokine staining and macrophage signature phenotypes, related to Figure 4**

(A) Single-plex chromogenic staining of IL6, IL8 & CXCL10 (IP10).

(B) Quantification of cytokine positive cell abundances.

(C) Bulk RNA levels of CXCL10 and IL8.

(D) PCA as in Fig. 4 K coloured by case.

(E) Volcano plots showing differential gene expression of each macrophage signature.

**Figure S5. Interferon alpha localisation, restriction factors and maternal decidual cytokine response, related to Figure 5**

(A) Interferon Alpha response gene expressions per NanoString GeoMx DSP compartment.

(B) Pathway analysis of significantly overexpressed genes in P^+^CoV19^CHI^ against P^-^CoV19^CHI^.

(C) Relationship between viral load and restriction factors for P^+^CoV19^CHI^ cases.

(D) Interferon blunting phenotype in macrophage compartments of P^+^CoV19^CHI^.

(E) IL6 and IL8 cytokine expression in maternal decidua by RNAscope.

**Figure S6. Virus microenvironment schematic, villous stroma response to SARS-CoV-2 and trophoblast signature phenotypes, related to Figure 6**

(A) Schematic of virus microenvironment analysis.

(B) Volcano plot of virus high vs virus low associated villous stroma.

(C) PCA as in Fig. 6 E coloured by SARS-CoV-2 postitivity and shaped by case.

(D) Volcano plots showing differential gene expression of each trophoblast signature.

(E) Schematic illustration of trophoblast signatures.

**Figure S7. Pathological and clinical signature of COVID-19 placentitis, timeline and gestation of disease and overview of study findings, related to Figure 7**

(A) Unbiased MCA of pathological and clinical data.

(B) Days from evidence of COVID-19 to IUD/birth.

(C) Gestation length of each COVID-19 associated disease group.

(D) Schematic illustration of overall study findings.

## Key Resources Table

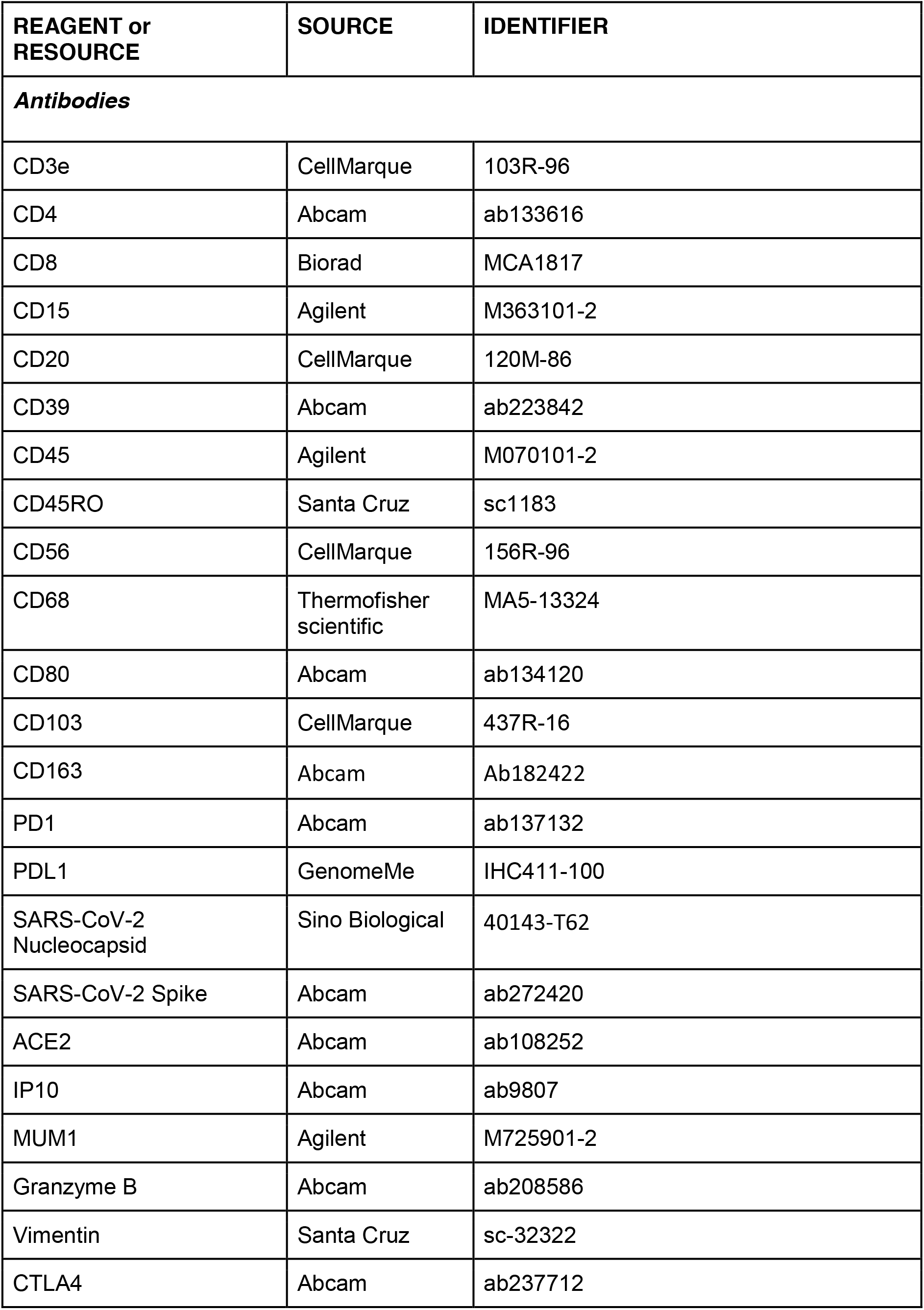
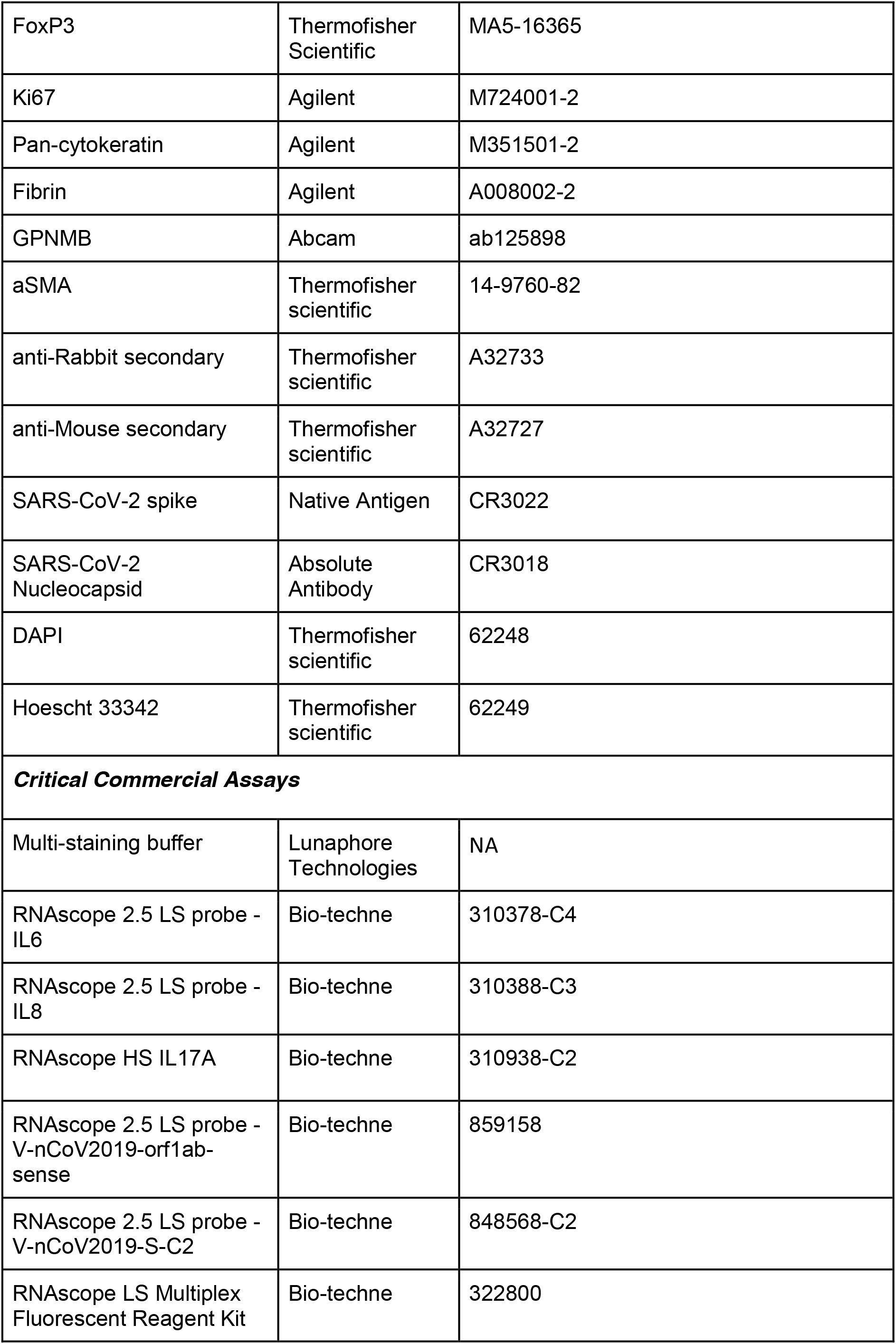
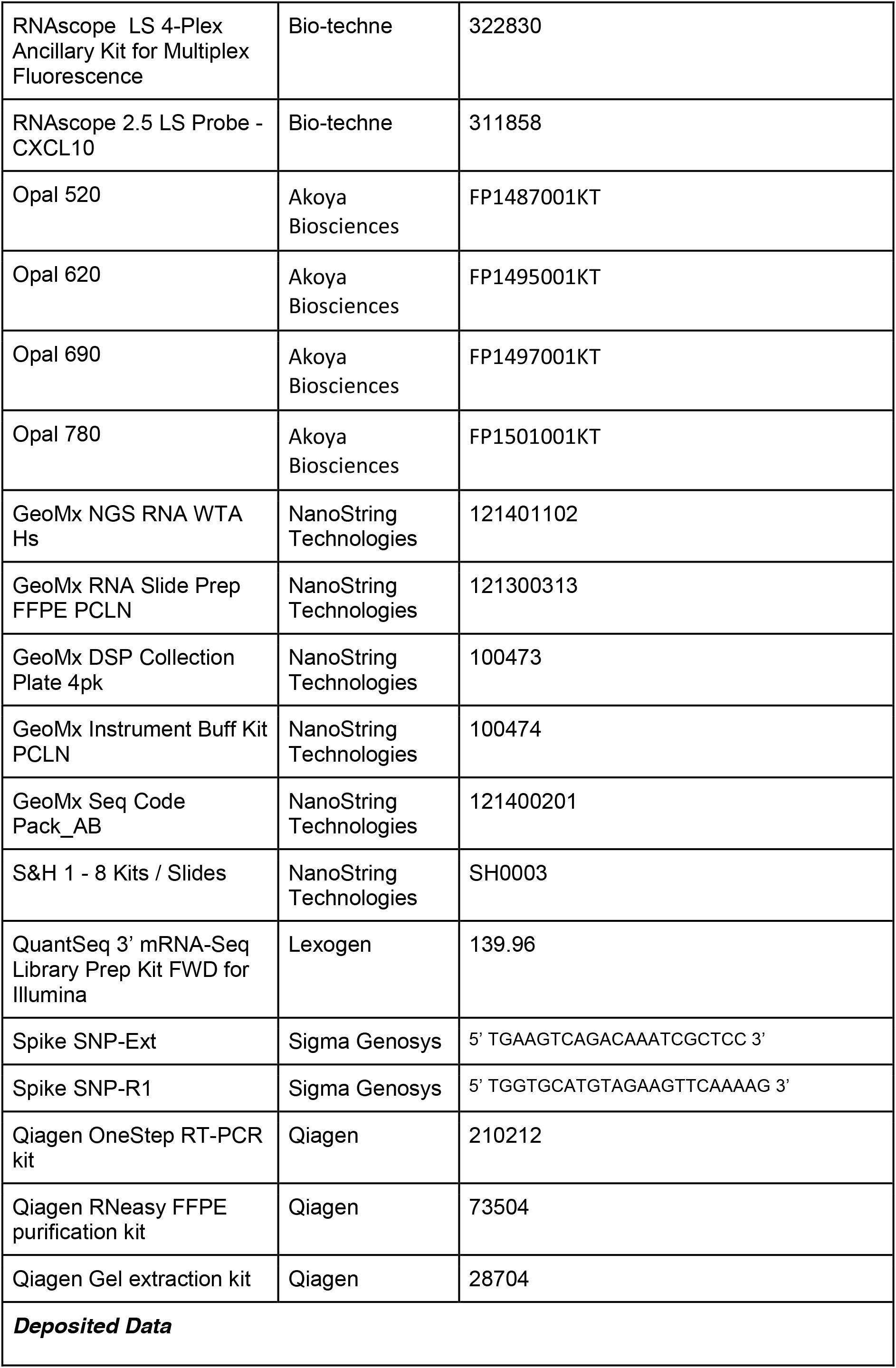
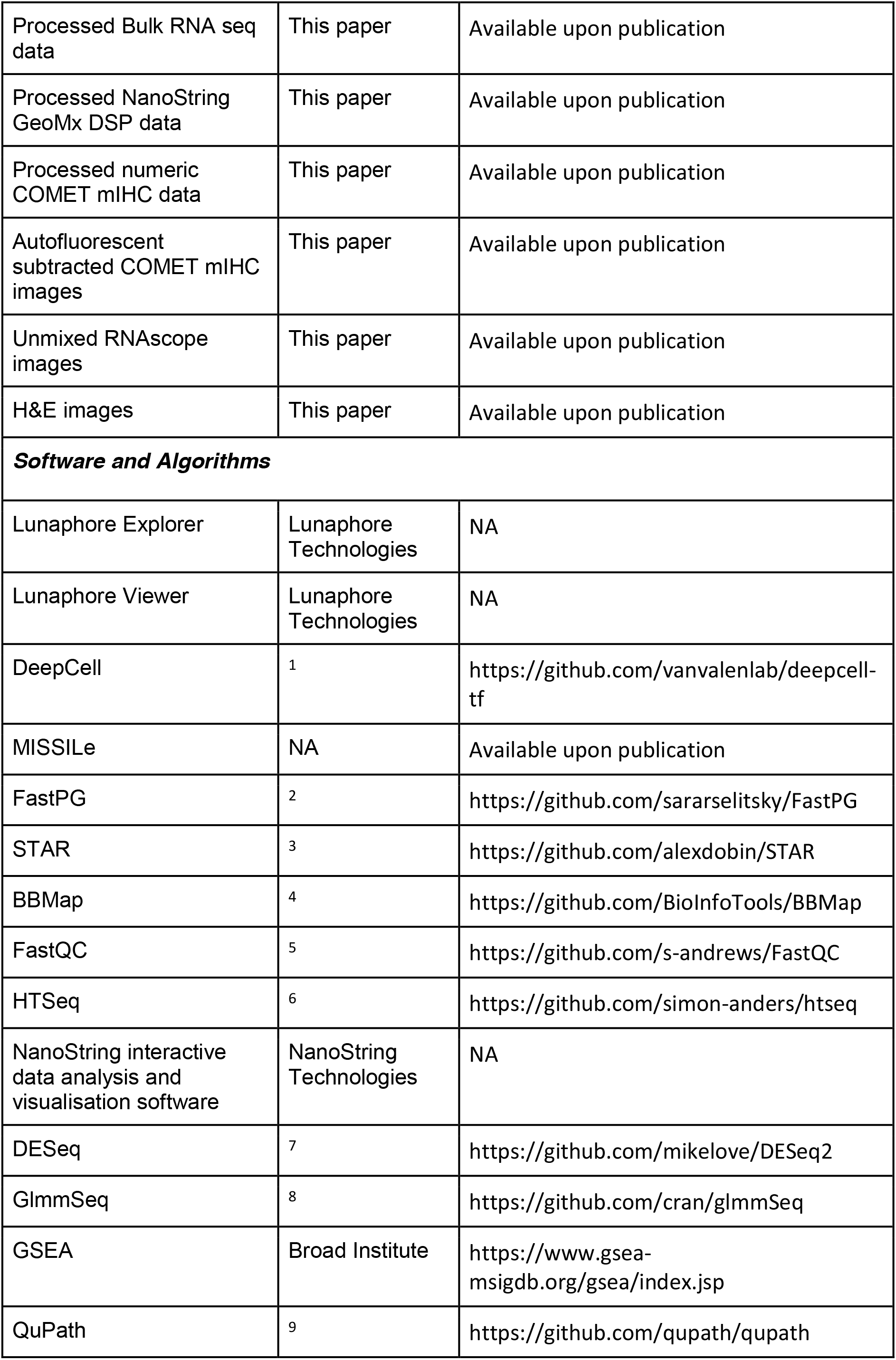
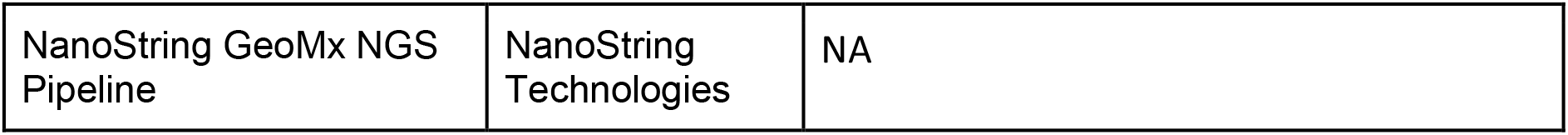

## References

1. Delorme-Axford, E., Sadovsky, Y., and Coyne, C.B. (2014). The Placenta as a Barrier to Viral Infections. Annu Rev Virol 1, 133–146. 10.1146/annurev-virology-031413-085524.

2. Hoo, R., Nakimuli, A., and Vento-Tormo, R. (2020). Innate Immune Mechanisms to Protect Against Infection at the Human Decidual-Placental Interface. Frontiers in Immunology 11. 10.3389/fimmu.2020.02070.

3. Wells, A.I., and Coyne, C.B. (2018). Type III Interferons in Antiviral Defenses at Barrier Surfaces. Trends Immunol 39, 848–858. 10.1016/j.it.2018.08.008.

4. Senegas, A., Villard, O., Neuville, A., Marcellin, L., Pfaff, A.W., Steinmetz, T., Mousli, M., Klein, J.P., and Candolfi, E. (2009). Toxoplasma gondii-induced foetal resorption in mice involves interferon-gamma-induced apoptosis and spiral artery dilation at the maternofoetal interface. Int J Parasitol 39, 481–487. 10.1016/j.ijpara.2008.08.009.

5. Yockey, L.J., Jurado, K.A., Arora, N., Millet, A., Rakib, T., Milano, K.M., Hastings, A.K., Fikrig, E., Kong, Y., Horvath, T.L., et al. (2018). Type I interferons instigate fetal demise after Zika virus infection. Sci Immunol 3. 10.1126/sciimmunol.aao1680.

6. Allotey, J., Stallings, E., Bonet, M., Yap, M., Chatterjee, S., Kew, T., Debenham, L., Llavall, A. C., Dixit, A., Zhou, D., et al. (2020). Clinical manifestations, risk factors, and maternal and perinatal outcomes of coronavirus disease 2019 in pregnancy: living systematic review and meta-analysis. Bmj 370, m3320. 10.1136/bmj.m3320.

7. Delahoy, M.J., Whitaker, M., O’Halloran, A., Chai, S.J., Kirley, P.D., Alden, N., Kawasaki, B., Meek, J., Yousey-Hindes, K., Anderson, E.J., et al. (2020). Characteristics and Maternal and Birth Outcomes of Hospitalized Pregnant Women with Laboratory-Confirmed COVID-19 - COVID-NET, 13 States, March 1-August 22, 2020. MMWR Morb Mortal Wkly Rep 69, 1347–1354. 10.15585/mmwr.mm6938e1.

8. Vousden, N., Ramakrishnan, R., Bunch, K., Morris, E., Simpson, N., Gale, C., O’Brien, P., Quigley, M., Brocklehurst, P., Kurinczuk, J.J., and Knight, M. (2022). Management and implications of severe COVID-19 in pregnancy in the UK: data from the UK Obstetric Surveillance System national cohort. Acta Obstet Gynecol Scand 101, 461–470. 10.1111/aogs.14329.

9. Babal, P., Krivosikova, L., Sarvaicova, L., Deckov, I., Szemes, T., Sedlackova, T., Palkovic, M., Kalinakova, A., and Janega, P. (2021). Intrauterine Fetal Demise After Uncomplicated COVID-19: What Can We Learn from the Case? Viruses 13. 10.3390/v13122545.

10. Schwartz, D.A., Avvad-Portari, E., Babál, P., Baldewijns, M., Blomberg, M., Bouachba, A., Camacho, J., Collardeau-Frachon, S., Colson, A., Dehaene, I., et al. (2022). Placental Tissue Destruction and Insufficiency from COVID-19 Causes Stillbirth and Neonatal Death from Hypoxic-Ischemic Injury: A Study of 68 Cases with SARS-CoV-2 Placentitis from 12 Countries. Archives of Pathology & Laboratory Medicine. 10.5858/arpa.2022-0029-SA.

11. Marton, T., Hargitai, B., Hunter, K., Pugh, M., and Murray, P. (2021). Massive Perivillous Fibrin Deposition and Chronic Histiocytic Intervillositis a Complication of SARS-CoV-2 Infection. Pediatr Dev Pathol 24, 450–454. 10.1177/10935266211020723.

12. Watkins, J.C., Torous, V.F., and Roberts, D.J. (2021). Defining Severe Acute Respiratory Syndrome Coronavirus 2 (SARS-CoV-2) Placentitis. Arch Pathol Lab Med 145, 1341–1349. 10.5858/arpa.2021-0246-SA.

13. Schwartz, D.A. (2020). An Analysis of 38 Pregnant Women With COVID-19, Their Newborn Infants, and Maternal-Fetal Transmission of SARS-CoV-2: Maternal Coronavirus Infections and Pregnancy Outcomes. Arch Pathol Lab Med 144, 799–805. 10.5858/arpa.2020-0901-SA.

14. Argueta, L.B., Lacko, L.A., Bram, Y., Tada, T., Carrau, L., Rendeiro, A.F., Zhang, T., Uhl, S., Lubor, B.C., Chandar, V., et al. (2022). Inflammatory responses in the placenta upon SARS-CoV-2 infection late in pregnancy. iScience 25, 104223. 10.1016/j.isci.2022.104223.

15. Linehan, L., O’Donoghue, K., Dineen, S., White, J., Higgins, J.R., and Fitzgerald, B. (2021). SARS-CoV-2 placentitis: An uncommon complication of maternal COVID-19. Placenta 104, 261–266. 10.1016/j.placenta.2021.01.012.

16. Stenton, S., McPartland, J., Shukla, R., Turner, K., Marton, T., Hargitai, B., Bamber, A., Pryce, J., Peres, C.L., Burguess, N., et al. (2022). SARS-COV2 placentitis and pregnancy outcome: A multicentre experience during the Alpha and early Delta waves of coronavirus pandemic in England. EClinicalMedicine 47, 101389. 10.1016/j.eclinm.2022.101389.

17. Desai, N., Neyaz, A., Szabolcs, A., Shih, A.R., Chen, J.H., Thapar, V., Nieman, L.T., Solovyov, A., Mehta, A., Lieb, D.J., et al. (2020). Temporal and spatial heterogeneity of host response to SARS-CoV-2 pulmonary infection. Nat Commun 11, 6319. 10.1038/s41467-020-20139-7.

18. Caniego-Casas, T., Martinez-Garcia, L., Alonso-Riano, M., Pizarro, D., Carretero-Barrio, I., Martinez-de-Castro, N., Ruz-Caracuel, I., de Pablo, R., Saiz, A., Royo, R.N., et al. (2022). RNA SARS-CoV-2 Persistence in the Lung of Severe COVID-19 Patients: A Case Series of Autopsies. Front Microbiol 13, 824967. 10.3389/fmicb.2022.824967.

19. Bodenheimer, T., Halappanavar, M., Jefferys, S., Gibson, R., Liu, S., Mucha, P.J., Stanley, N., Parker, J.S., and Selitsky, S.R. (2020). FastPG: fast clustering of millions of single cells. BioRxiv.

20. Braun, E., Hotter, D., Koepke, L., Zech, F., Groß, R., Sparrer, K.M.J., Müller, J.A., Pfaller, C.K., Heusinger, E., Wombacher, R., et al. (2019). Guanylate-Binding Proteins 2 and 5 Exert Broad Antiviral Activity by Inhibiting Furin-Mediated Processing of Viral Envelope Proteins. Cell Rep 27, 2092–2104.e2010. 10.1016/j.celrep.2019.04.063.

21. Schenkel, J.M., and Masopust, D. (2014). Tissue-resident memory T cells. Immunity 41, 886–897. 10.1016/j.immuni.2014.12.007.

22. Merad, M., and Martin, J.C. (2020). Pathological inflammation in patients with COVID-19: a key role for monocytes and macrophages. Nat Rev Immunol 20, 355–362. 10.1038/s41577-020-0331-4.

23. Santa Cruz, A., Mendes-Frias, A., Oliveira, A.I., Dias, L., Matos, A.R., Carvalho, A., Capela, C., Pedrosa, J., Castro, A.G., and Silvestre, R. (2021). Interleukin-6 Is a Biomarker for the Development of Fatal Severe Acute Respiratory Syndrome Coronavirus 2 Pneumonia. Front Immunol 12, 613422. 10.3389/fimmu.2021.613422.

24. Li, L., Li, J., Gao, M., Fan, H., Wang, Y., Xu, X., Chen, C., Liu, J., Kim, J., Aliyari, R., et al. (2020). Interleukin-8 as a Biomarker for Disease Prognosis of Coronavirus Disease-2019 Patients. Front Immunol 11, 602395. 10.3389/fimmu.2020.602395.

25. Zhang, N., Zhao, Y.D., and Wang, X.M. (2020). CXCL10 an important chemokine associated with cytokine storm in COVID-19 infected patients. Eur Rev Med Pharmacol Sci 24, 7497–7505. 10.26355/eurrev_202007_21922.

26. Chidambaranathan-Reghupaty, S., Mendoza, R., Fisher, P.B., and Sarkar, D. (2018). The multifaceted oncogene SND1 in cancer: focus on hepatocellular carcinoma. Hepatoma Res 4. 10.20517/2394-5079.2018.34.

27. Tsukamoto, Y., Helsel, W.E., and Wahl, S.M. (1981). Macrophage production of fibronectin, a chemoattractant for fibroblasts. J Immunol 127, 673–678.

28. Duan, Y., Zhang, L., Angosto-Bazarra, D., Pelegrin, P., Nunez, G., and He, Y. (2020). RACK1 Mediates NLRP3 Inflammasome Activation by Promoting NLRP3 Active Conformation and Inflammasome Assembly. Cell Rep 33, 108405. 10.1016/j.celrep.2020.108405.

29. Jiang, Z., Chen, Z., Hu, L., Qiu, L., and Zhu, L. (2020). Calreticulin Blockade Attenuates Murine Acute Lung Injury by Inducing Polarization of M2 Subtype Macrophages. Front Immunol 11, 11. 10.3389/fimmu.2020.00011.

30. Martin-Sancho, L., Lewinski, M.K., Pache, L., Stoneham, C.A., Yin, X., Becker, M.E., Pratt, D., Churas, C., Rosenthal, S.B., and Liu, S. (2021). Functional landscape of SARS-CoV-2 cellular restriction. Molecular cell 81, 2656–2668. e2658.

31. Zhou, Z., Ren, L., Zhang, L., Zhong, J., Xiao, Y., Jia, Z., Guo, L., Yang, J., Wang, C., Jiang, S., et al. (2020). Heightened Innate Immune Responses in the Respiratory Tract of COVID-19 Patients. Cell Host Microbe 27, 883–890 e882. 10.1016/j.chom.2020.04.017.

32. Winstone, H., Lista, M.J., Reid, A.C., Bouton, C., Pickering, S., Galao, R.P., Kerridge, C., Doores, K.J., Swanson, C.M., and Neil, S.J.D. (2021). The Polybasic Cleavage Site in SARS-CoV-2 Spike Modulates Viral Sensitivity to Type I Interferon and IFITM2. J Virol 95. 10.1128/JVI.02422-20.

33. Anderson, S.L., Carton, J.M., Lou, J., Xing, L., and Rubin, B.Y. (1999). Interferon-induced guanylate binding protein-1 (GBP-1) mediates an antiviral effect against vesicular stomatitis virus and encephalomyocarditis virus. Virology 256, 8–14. 10.1006/viro.1999.9614.

34. Lokugamage, K.G., Hage, A., de Vries, M., Valero-Jimenez, A.M., Schindewolf, C., Dittmann, M., Rajsbaum, R., and Menachery, V.D. (2020). Type I Interferon Susceptibility Distinguishes SARS-CoV-2 from SARS-CoV. J Virol 94. 10.1128/JVI.01410-20.

35. Chua, R.L., Lukassen, S., Trump, S., Hennig, B.P., Wendisch, D., Pott, F., Debnath, O., Thurmann, L., Kurth, F., Volker, M.T., et al. (2020). COVID-19 severity correlates with airway epithelium-immune cell interactions identified by single-cell analysis. Nat Biotechnol 38, 970–979. 10.1038/s41587-020-0602-4.

36. Stone, R.C., Pastar, I., Ojeh, N., Chen, V., Liu, S., Garzon, K.I., and Tomic-Canic, M. (2016). Epithelial-mesenchymal transition in tissue repair and fibrosis. Cell Tissue Res 365, 495–506. 10.1007/s00441-016-2464-0.

37. Stewart, C.A., Gay, C.M., Ramkumar, K., Cargill, K.R., Cardnell, R.J., Nilsson, M.B., Heeke, S., Park, E.M., Kundu, S.T., Diao, L., et al. (2021). Lung Cancer Models Reveal Severe Acute Respiratory Syndrome Coronavirus 2-Induced Epithelial-to-Mesenchymal Transition Contributes to Coronavirus Disease 2019 Pathophysiology. J Thorac Oncol 16, 1821–1839. 10.1016/j.jtho.2021.07.002.

38. Mathew, D., Giles Josephine, R., Baxter Amy, E., Oldridge Derek, A., Greenplate Allison, R., Wu Jennifer, E., Alanio, C., Kuri-Cervantes, L., Pampena, M.B., D’Andrea, K., et al. (2020). Deep immune profiling of COVID-19 patients reveals distinct immunotypes with therapeutic implications. Science 369, eabc8511. 10.1126/science.abc8511.

39. Sefik, E., Qu, R., Junqueira, C., Kaffe, E., Mirza, H., Zhao, J., Brewer, J.R., Han, A., Steach, H.R., Israelow, B., et al. (2022). Inflammasome activation in infected macrophages drives COVID-19 pathology. Nature 606, 585–593. 10.1038/s41586-022-04802-1.

40. Delorey, T.M., Ziegler, C.G.K., Heimberg, G., Normand, R., Yang, Y., Segerstolpe, A., Abbondanza, D., Fleming, S.J., Subramanian, A., Montoro, D.T., et al. (2021). COVID-19 tissue atlases reveal SARS-CoV-2 pathology and cellular targets. Nature 595, 107–113. 10.1038/s41586-021-03570-8.

41. Melms, J.C., Biermann, J., Huang, H., Wang, Y., Nair, A., Tagore, S., Katsyv, I., Rendeiro, A.F., Amin, A.D., Schapiro, D., et al. (2021). A molecular single-cell lung atlas of lethal COVID-19. Nature 595, 114–119. 10.1038/s41586-021-03569-1.

42. Fahmi, A., Brugger, M., Demoulins, T., Zumkehr, B., Oliveira Esteves, B.I., Bracher, L., Wotzkow, C., Blank, F., Thiel, V., Baud, D., and Alves, M.P. (2021). SARS-CoV-2 can infect and propagate in human placenta explants. Cell Rep Med 2, 100456. 10.1016/j.xcrm.2021.100456.

43. Brady, C.A., Williams, C., Sharps, M.C., Shelleh, A., Batra, G., Heazell, A.E.P., and Crocker, I.P. (2021). Chronic histiocytic intervillositis: A breakdown in immune tolerance comparable to allograft rejection? Am J Reprod Immunol 85, e13373. 10.1111/aji.13373.

44. Zhang, F., Mears, J.R., Shakib, L., Beynor, J.I., Shanaj, S., Korsunsky, I., Nathan, A., Accelerating Medicines Partnership Rheumatoid, A., Systemic Lupus Erythematosus, C., Donlin, L.T., and Raychaudhuri, S. (2021). IFN-gamma and TNF-alpha drive a CXCL10+ CCL2+ macrophage phenotype expanded in severe COVID-19 lungs and inflammatory diseases with tissue inflammation. Genome Med 13, 64. 10.1186/s13073-021-00881-3.

45. Lin, T., Geng, T., Harrison, A.G., Yang, D., Vella, A.T., Fikrig, E., and Wang, P. (2020). CXCL10 Signaling Contributes to the Pathogenesis of Arthritogenic Alphaviruses. Viruses 12. 10.3390/v12111252.

46. Mauricio, R., and Khera, A. (2022). Statin Use in Pregnancy: Is It Time For a Paradigm Shift? Circulation 145, 496–498. 10.1161/CIRCULATIONAHA.121.058983.

47. Gudowska-Sawczuk, M., and Mroczko, B. (2022). What Is Currently Known about the Role of CXCL10 in SARS-CoV-2 Infection? Int J Mol Sci 23. 10.3390/ijms23073673.

48. Thomas, J., Sun, Y., and Debelenko, L. (2021). Infrequent Placental and Fetal Involvement in SARS-CoV-2 Infection: Pathology Data from a Large Medical Center. J Dev Biol 9. 10.3390/jdb9040045.

49. Hikmet, F., Mear, L., Edvinsson, A., Micke, P., Uhlen, M., and Lindskog, C. (2020). The protein expression profile of ACE2 in human tissues. Mol Syst Biol 16, e9610. 10.15252/msb.20209610.

50. Hecht, J.L., Quade, B., Deshpande, V., Mino-Kenudson, M., Ting, D.T., Desai, N., Dygulska, B., Heyman, T., Salafia, C., Shen, D., et al. (2020). SARS-CoV-2 can infect the placenta and is not associated with specific placental histopathology: a series of 19 placentas from COVID-19-positive mothers. Mod Pathol 33, 2092–2103. 10.1038/s41379-020-0639-4.

51. Edlow, A.G., Li, J.Z., Collier, A.Y., Atyeo, C., James, K.E., Boatin, A.A., Gray, K.J., Bordt, E.A., Shook, L.L., Yonker, L.M., et al. (2020). Assessment of Maternal and Neonatal SARS-CoV-2 Viral Load, Transplacental Antibody Transfer, and Placental Pathology in Pregnancies During the COVID-19 Pandemic. JAMA Netw Open 3, e2030455. 10.1001/jamanetworkopen.2020.30455.

52. Zimmerman, S.M., Fropf, R., Kulasekara, B.R., Griswold, M., Appelbe, O., Bahrami, A., Boykin, R., Buhr, D.L., Fuhrman, K., Hoang, M.L., et al. (2022). Spatially resolved whole transcriptome profiling in human and mouse tissue using Digital Spatial Profiling. Genome Res. 10.1101/gr.276206.121.

53. Babiker, A., Immergluck, K., Stampfer, S.D., Rao, A., Bassit, L., Su, M., Nguyen, V., Stittleburg, V., Ingersoll, J.M., Bradley, H.L., et al. (2021). Single-Amplicon Multiplex Real-Time Reverse Transcription-PCR with Tiled Probes To Detect SARS-CoV-2 spike Mutations Associated with Variants of Concern. J Clin Microbiol 59, e0144621. 10.1128/JCM.01446-21.

54. Greenwald, N.F., Miller, G., Moen, E., Kong, A., Kagel, A., Dougherty, T., Fullaway, C.C., McIntosh, B.J., Leow, K.X., Schwartz, M.S., et al. (2022). Whole-cell segmentation of tissue images with human-level performance using large-scale data annotation and deep learning. Nat Biotechnol 40, 555–565. 10.1038/s41587-021-01094-0.

55. Schurch, C.M., Bhate, S.S., Barlow, G.L., Phillips, D.J., Noti, L., Zlobec, I., Chu, P., Black, S., Demeter, J., McIlwain, D.R., et al. (2020). Coordinated Cellular Neighborhoods Orchestrate Antitumoral Immunity at the Colorectal Cancer Invasive Front. Cell 182, 1341–1359 e1319. 10.1016/j.cell.2020.07.005.

56. Jiang, S., Chan, C.N., Rovira-Clave, X., Chen, H., Bai, Y., Zhu, B., McCaffrey, E., Greenwald, N.F., Liu, C., Barlow, G.L., et al. (2022). Combined protein and nucleic acid imaging reveals virus-dependent B cell and macrophage immunosuppression of tissue microenvironments. Immunity 55, 1118–1134 e1118. 10.1016/j.immuni.2022.03.020.

57. Bankhead, P., Loughrey, M.B., Fernandez, J.A., Dombrowski, Y., McArt, D.G., Dunne, P.D., McQuaid, S., Gray, R.T., Murray, L.J., Coleman, H.G., et al. (2017). QuPath: Open source software for digital pathology image analysis. Sci Rep 7, 16878. 10.1038/s41598-017-17204-5.

58. Myles Lewi, K.G., Elisabetta Sciacca, Cankut Cubuk, Anna Surace (2022). glmmSeq: General Linear Mixed Models for Gene-level Differential Expression. https://github.com/myles-lewis/glmmSeq.

59. Andrews, S. (2010). FastQC: A Quality Control Tool for High Throughput Sequence Data.

60. Dobin, A., Davis, C.A., Schlesinger, F., Drenkow, J., Zaleski, C., Jha, S., Batut, P., Chaisson, M., and Gingeras, T.R. (2013). STAR: ultrafast universal RNA-seq aligner. Bioinformatics 29, 15–21. 10.1093/bioinformatics/bts635.

61. Li, H., Handsaker, B., Wysoker, A., Fennell, T., Ruan, J., Homer, N., Marth, G., Abecasis, G., Durbin, R., and Genome Project Data Processing, S. (2009). The Sequence Alignment/Map format and SAMtools. Bioinformatics 25, 2078–2079. 10.1093/bioinformatics/btp352.

62. Putri, G.H., Anders, S., Pyl, P.T., Pimanda, J.E., and Zanini, F. (2022). Analysing high-throughput sequencing data in Python with HTSeq 2.0. Bioinformatics 38, 2943–2945. 10.1093/bioinformatics/btac166.

63. Love, M.I., Huber, W., and Anders, S. (2014). Moderated estimation of fold change and dispersion for RNA-seq data with DESeq2. Genome Biol 15, 550. 10.1186/s13059-014-0550-8.

64. Mootha, V.K., Lindgren, C.M., Eriksson, K.F., Subramanian, A., Sihag, S., Lehar, J., Puigserver, P., Carlsson, E., Ridderstrale, M., Laurila, E., et al. (2003). PGC-1alpha-responsive genes involved in oxidative phosphorylation are coordinately downregulated in human diabetes. Nat Genet 34, 267–273. 10.1038/ng1180.

65. Subramanian, A., Tamayo, P., Mootha, V.K., Mukherjee, S., Ebert, B.L., Gillette, M.A., Paulovich, A., Pomeroy, S.L., Golub, T.R., Lander, E.S., and Mesirov, J.P. (2005). Gene set enrichment analysis: a knowledge-based approach for interpreting genome-wide expression profiles. Proc Natl Acad Sci U S A 102, 15545–15550. 10.1073/pnas.0506580102.

66. Gu, Z., Eils, R., and Schlesner, M. (2016). Complex heatmaps reveal patterns and correlations in multidimensional genomic data. Bioinformatics 32, 2847–2849. 10.1093/bioinformatics/btw313.

## Key Resource Table References

1. Greenwald, N.F., Miller, G., Moen, E., Kong, A., Kagel, A., Dougherty, T., Fullaway, C.C., McIntosh, B.J., Leow, K.X., Schwartz, M.S., et al. (2022). Whole-cell segmentation of tissue images with human-level performance using large-scale data annotation and deep learning. Nat Biotechnol 40, 555–565. 10.1038/s41587-021-01094-0.

2. Bodenheimer, T., Halappanavar, M., Jefferys, S., Gibson, R., Liu, S., Mucha, P.J., Stanley, N., Parker, J.S., and Selitsky, S.R. (2020). FastPG: fast clustering of millions of single cells. BioRxiv.

3. Dobin, A., Davis, C.A., Schlesinger, F., Drenkow, J., Zaleski, C., Jha, S., Batut, P., Chaisson, M., and Gingeras, T.R. (2013). STAR: ultrafast universal RNA-seq aligner. Bioinformatics 29, 15–21. 10.1093/bioinformatics/bts635.

4. Bushnell, B. (2014). BBMap: A Fast, Accurate, Splice-Aware Aligner. LBNL Report.

5. Andrews, S. (2010). FastQC: A Quality Control Tool for High Throughput Sequence Data.

6. Putri, G.H., Anders, S., Pyl, P.T., Pimanda, J.E., and Zanini, F. (2022). Analysing high-throughput sequencing data in Python with HTSeq 2.0. Bioinformatics 38, 2943–2945. 10.1093/bioinformatics/btac166.

7. Love, M.I., Huber, W., and Anders, S. (2014). Moderated estimation of fold change and dispersion for RNA-seq data with DESeq2. Genome Biol 15, 550. 10.1186/s13059-014-0550-8.

8. Myles Lewi, K.G., Elisabetta Sciacca, Cankut Cubuk, Anna Surace (2022). glmmSeq: General Linear Mixed Models for Gene-level Differential Expression. https://github.com/myles-lewis/glmmSeq.

9. Bankhead, P., Loughrey, M.B., Fernandez, J.A., Dombrowski, Y., McArt, D.G., Dunne, P.D., McQuaid, S., Gray, R.T., Murray, L.J., Coleman, H.G., et al. (2017). QuPath: Open source software for digital pathology image analysis. Sci Rep 7, 16878. 10.1038/s41598-017-17204-5.

