## Supplementary Tables and Figures for "Multi-omic spatial profiling reveals the unique virus-driven immune landscape of COVID-19 placentitis"

Figure S1

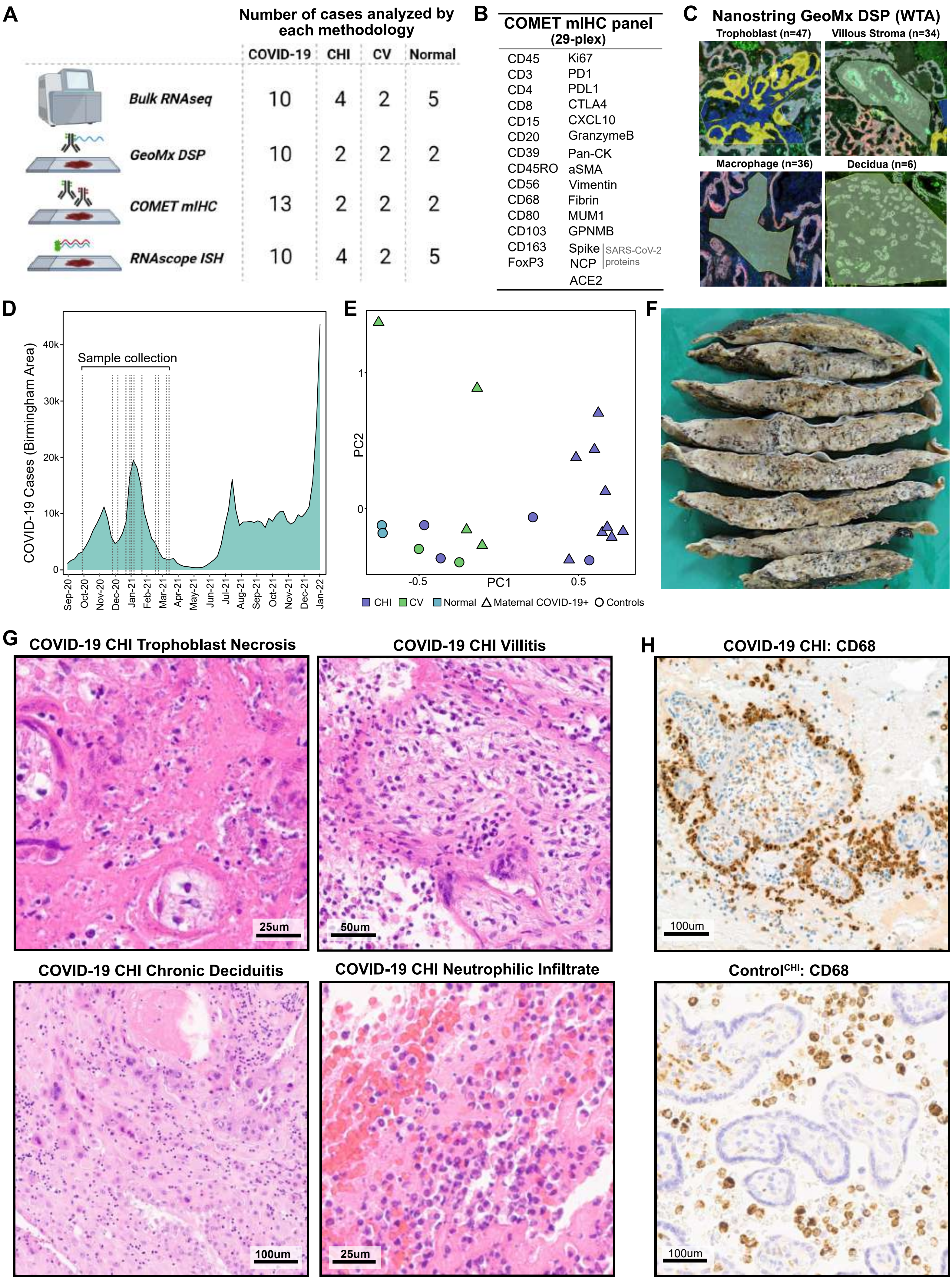
A
Number of cases analyzed by each methodology
COVID-19
CHI
CV
Normal
Bulk RNAseq
10
4
2
5
GeoMx DSP
10
2
2
2
COMET mIHC
13
2
2
2
RNAscope ISH
10
4
2
5
B
COMET mIHC panel (29-plex)
CD45 Ki67
CD3 PD1
CD4 PDL1
CD8 CTLA4
CD15 CXCL10
CD20 GranzymeB
CD39 Pan-CK
CD45RO aSMA
CD56 Vimentin
CD68 Fibrin
CD80 MUM1
CD103 GPNMB
CD163 Spike
FoxP3 NCP
SARS-CoV-2 proteins
ACE2
C
Nanostring GeoMx DSP (WTA)
Trophoblast (n=47)
Villous Stroma (n=34)
Macrophage (n=36)
Decidua (n=6)
D
Sample collection
COVID-19 Cases (Birmingham Area)
0
10k
20k
30k
40k
Sep-20 Oct-20 Nov-20 Dec-20 Jan-21 Feb-21 Mar-21 Apr-21 May-21 Jun-21 Jul-21 Aug-21 Sep-21 Oct-21 Nov-21 Dec-21 Jan-22
E
PC2
PC1
1
0
-0.5
0.5
CHI
CV
Normal
Maternal COVID-19+
Controls
F
G
COVID-19 CHI Trophoblast Necrosis
25um
COVID-19 CHI Villitis
50um
COVID-19 CHI Chronic Deciduitis
100um
COVID-19 CHI Neutrophilic Infiltrate
25um
H
COVID-19 CHI: CD68
100um
ControlCHI: CD68
100um

Figure S2

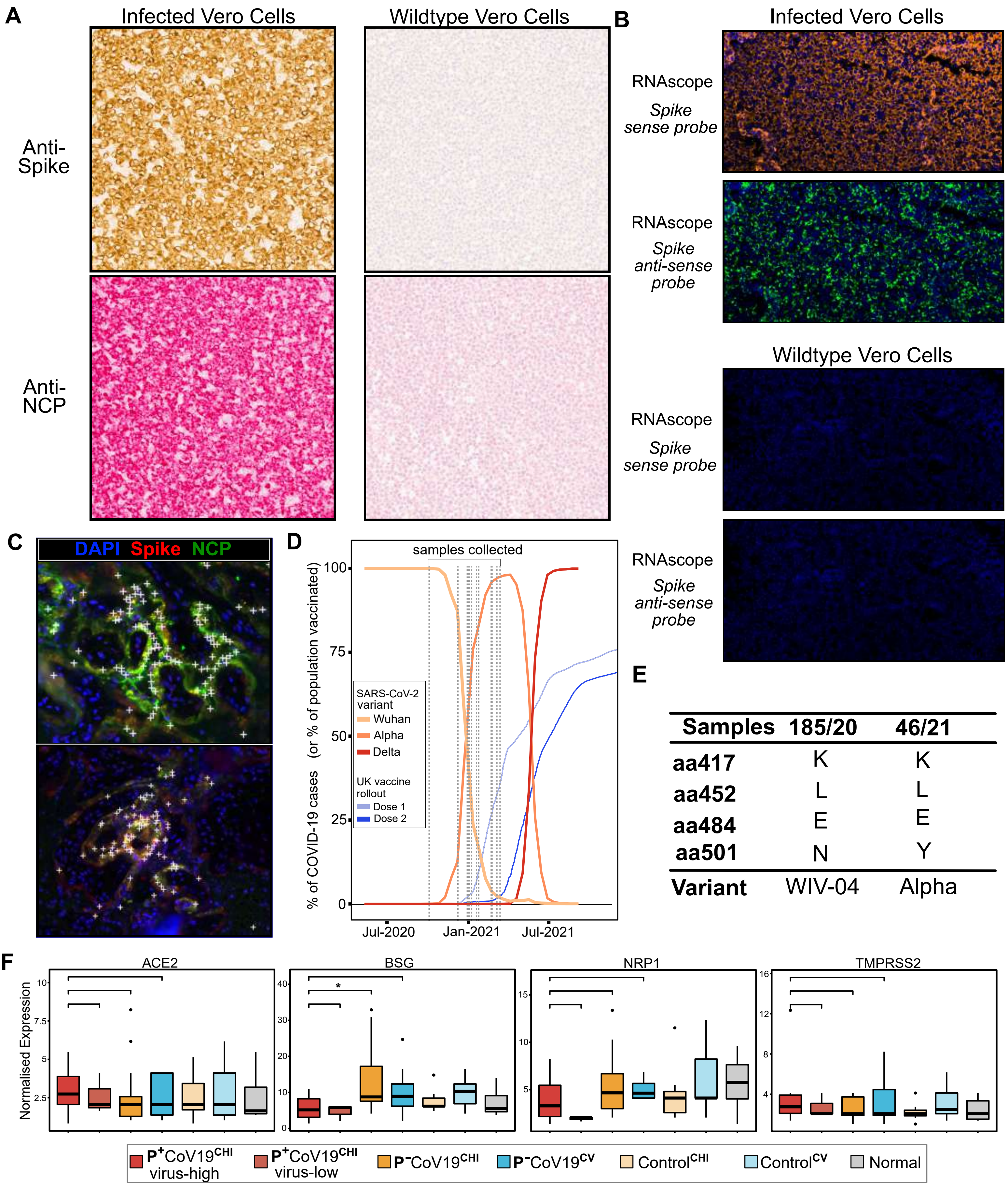
A
Infected Vero Cells
Wildtype Vero Cells
Anti-Spike
Anti-NCP
B
Infected Vero Cells
RNAscope Spike sense probe
RNAscope Spike anti-sense probe
Wildtype Vero Cells
RNAscope Spike sense probe
RNAscope Spike anti-sense probe
C
DAPI Spike NCP
D
samples collected
% of COVID-19 cases (or % of population vaccinated)
100
75
50
25
0
SARS-CoV-2 variant
Wuhan
Alpha
Delta
UK vaccine rollout
Dose 1
Dose 2
Jul-2020
Jan-2021
Jul-2021
E
Samples 185/20 46/21
aa417 K K
aa452 L L
aa484 E E
aa501 N Y
Variant WIV-04 Alpha
F
ACE2
BSG
NRP1
TMPRSS2
Normalised Expression
*
P+CoV19CHI virus-high
P+CoV19CHI virus-low
P-CoV19CHI
P-CoV19CV
ControlCHI
ControlCV
Normal

Figure S3

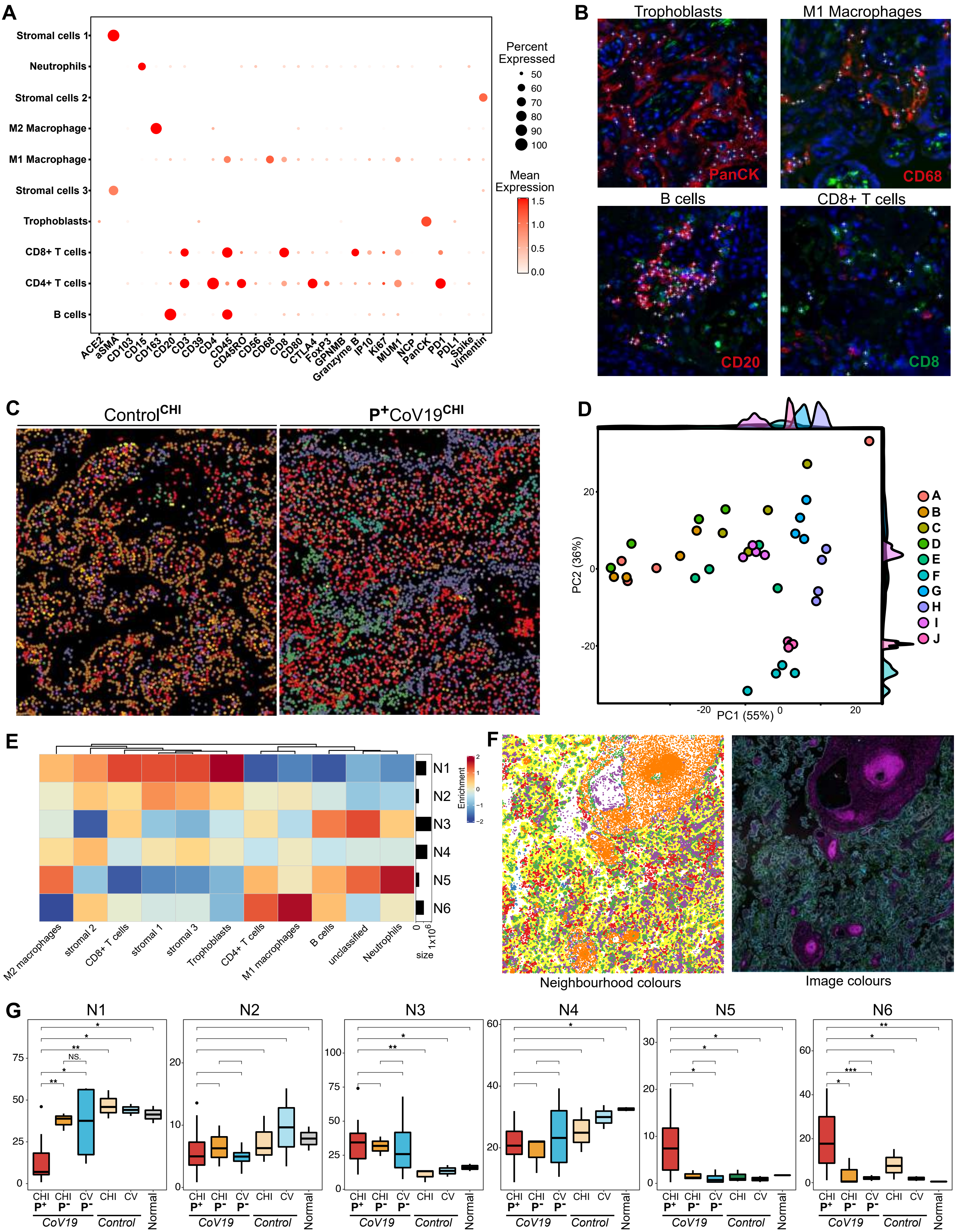

A
Stromal cells 1
Neutrophils
Stromal cells 2
M2 Macrophage
M1 Macrophage
Stromal cells 3
Trophoblasts
CD8+ T cells
CD4+ T cells
B cells
ACE2 aSMA CD103 CD15 CD163 CD20 CD3 CD39 CD4 CD45 CD45RO CD56 CD68 CD8 CD80 CTLA4 FoxP3 GPNMB Granzyme B IP10 Ki67 MUM1 NCP PanCK PD1 PDL1 Spike Vimentin
Percent Expressed
50 60 70 80 90 100
Mean Expression
1.5 1.0 0.5 0.0
B
Trophoblasts
PanCK
M1 Macrophages
CD68
B cells
CD20
CD8+ T cells
CD8
C
Control CHI
P+CoV19 CHI
D
PC1 (55%)
PC2 (36%)
A B C D E F G H I J
E
N1 N2 N3 N4 N5 N6
Enrichment
M2 macrophages stromal 2 CD8+ T cells stromal 1 stromal 3 Trophoblasts CD4+ T cells M1 macrophages B cells unclassified Neutrophils
size
F
Neighbourhood colours
Image colours
G
N1 N2 N3 N4 N5 N6
CHI P+ CHI P- CV P- CHI CV
CoV19 Control Normal

### Figure S4

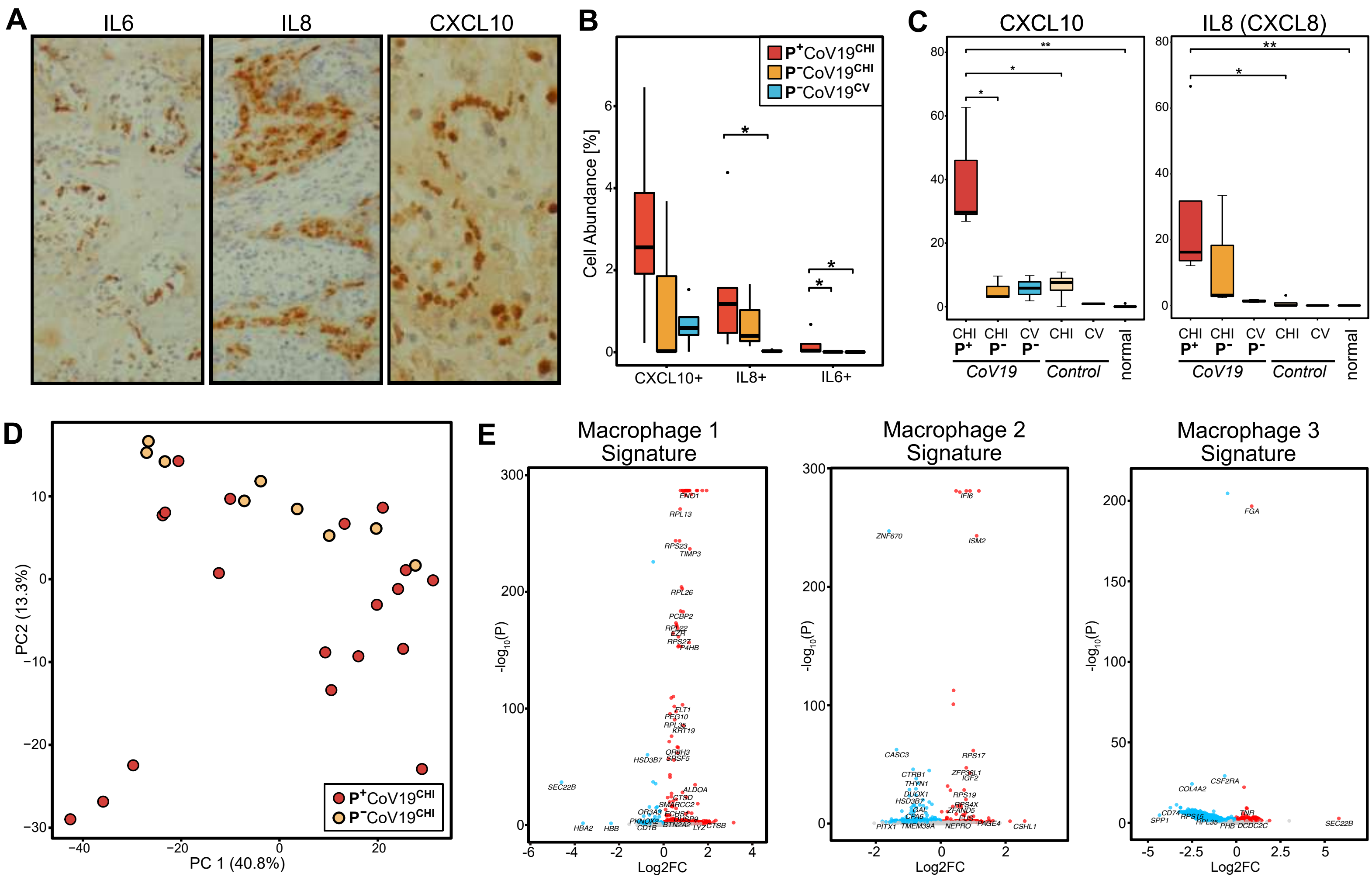
A
IL6
IL8
CXCL10
B
P+CoV19CHI
P-CoV19CHI
P-CoV19CV
Cell Abundance [%]
CXCL10+
IL8+
IL6+
C
CXCL10
IL8 (CXCL8)
CHI P+
CHI P-
CV P-
CHI
CV
CoV19
Control
normal
D
PC2 (13.3%)
PC 1 (40.8%)
P+CoV19CHI
P-CoV19CHI
E
Macrophage 1 Signature
Macrophage 2 Signature
Macrophage 3 Signature
-log10(P)
Log2FC

Figure S5

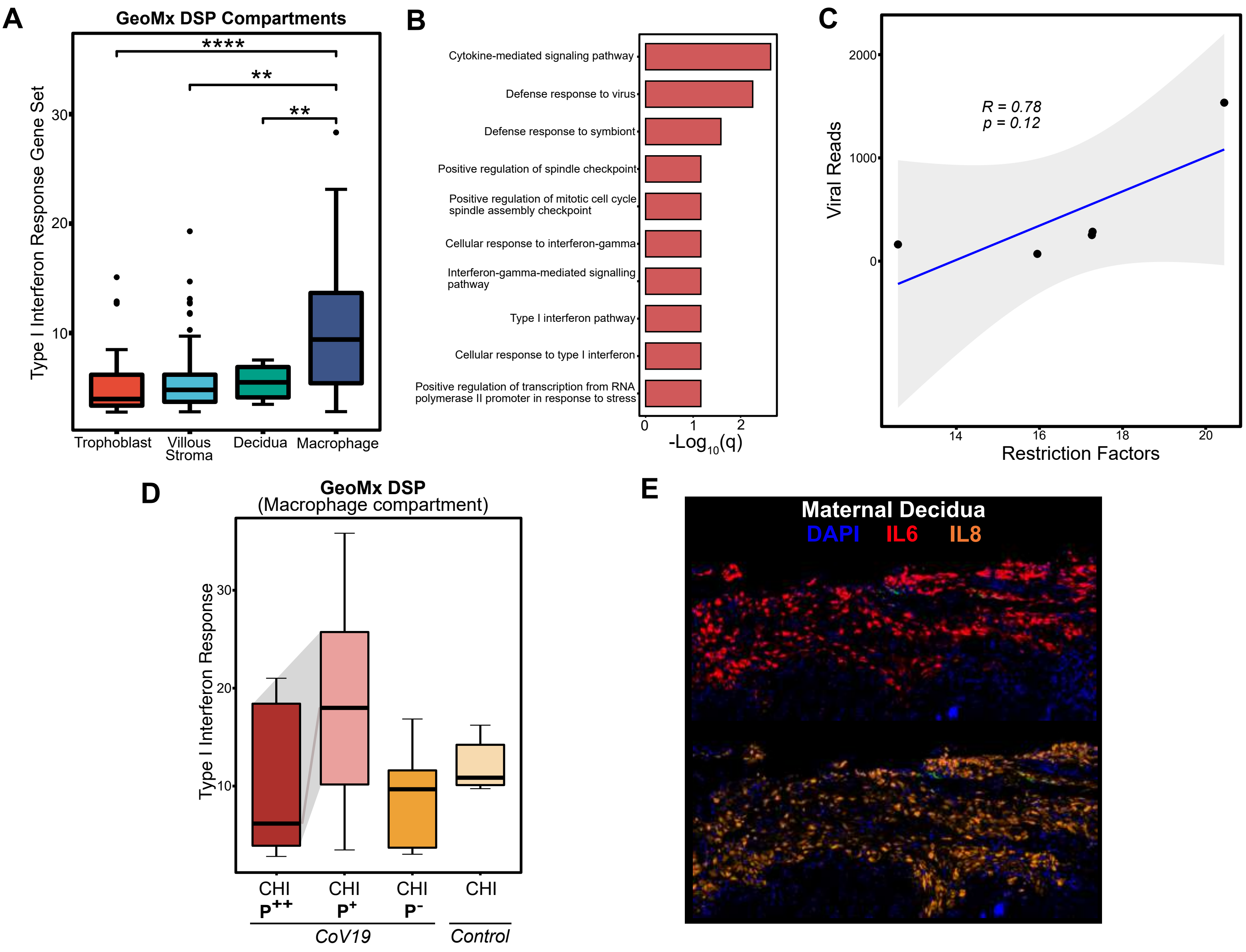
A
GeoMx DSP Compartments
Type I Interferon Response Gene Set
Trophoblast
Villous Stroma
Decidua
Macrophage
B
Cytokine-mediated signaling pathway
Defense response to virus
Defense response to symbiont
Positive regulation of spindle checkpoint
Positive regulation of mitotic cell cycle spindle assembly checkpoint
Cellular response to interferon-gamma
Interferon-gamma-mediated signalling pathway
Type I interferon pathway
Cellular response to type I interferon
Positive regulation of transcription from RNA polymerase II promoter in response to stress
-Log10(q)
C
R = 0.78
p = 0.12
Viral Reads
Restriction Factors
D
GeoMx DSP
(Macrophage compartment)
Type I Interferon Response
CHI P++
CHI P+
CHI P-
CHI
CoV19
Control
E
Maternal Decidua
DAPI
IL6
IL8

Figure S6

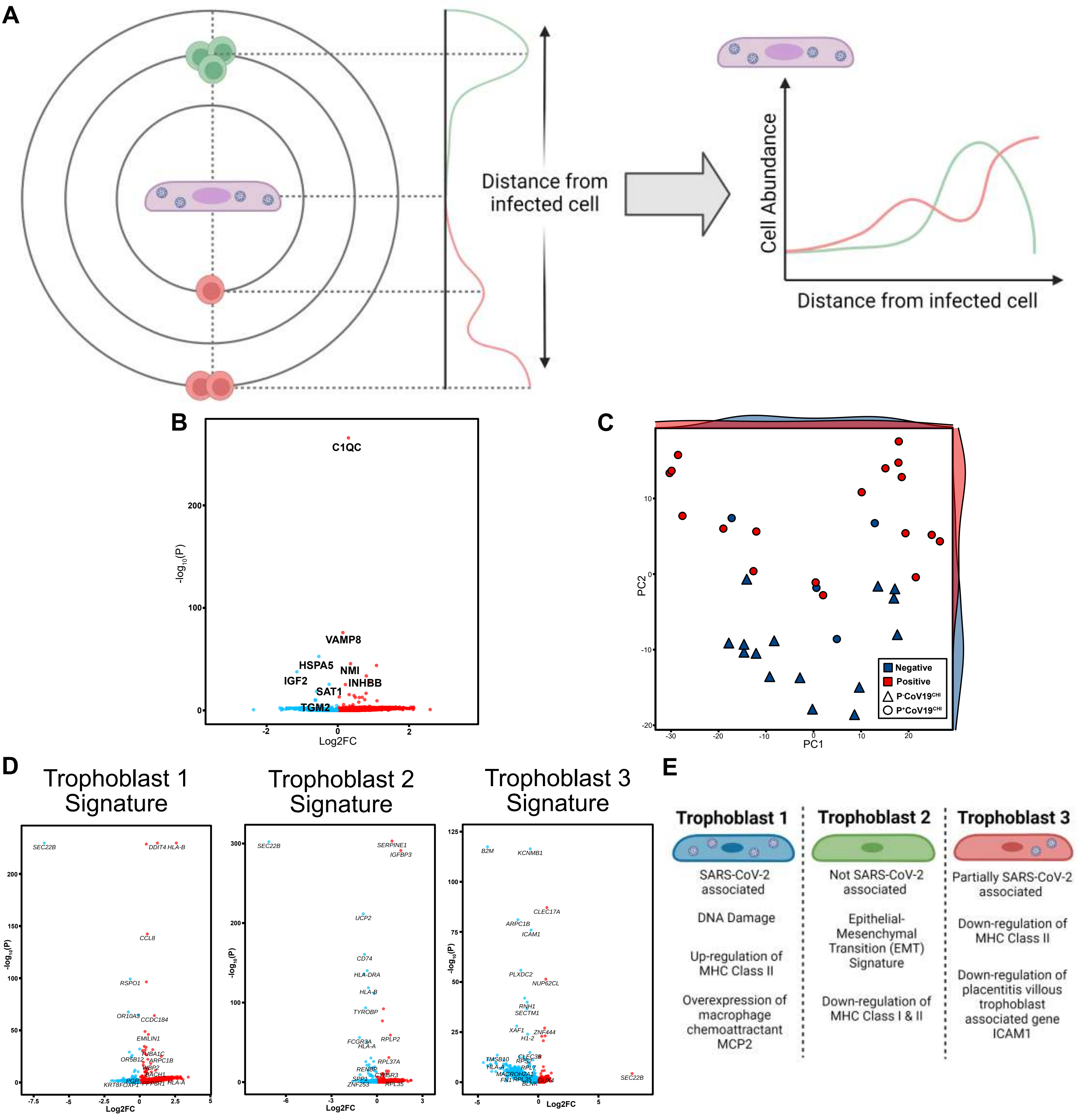

A
Distance from infected cell
Cell Abundance
Distance from infected cell
B
C1QC
VAMP8
HSPA5
IGF2
NMI
INHBB
SAT1
TGM2
-log10(P)
Log2FC
C
PC1
PC2
Negative
Positive
P-CoV19CHI
P+CoV19CHI
D
Trophoblast 1 Signature
Trophoblast 2 Signature
Trophoblast 3 Signature
-log10(P)
Log2FC
E
Trophoblast 1
SARS-CoV-2 associated
DNA Damage
Up-regulation of MHC Class II
Overexpression of macrophage chemoattractant MCP2
Trophoblast 2
Not SARS-CoV-2 associated
Epithelial-Mesenchymal Transition (EMT) Signature
Down-regulation of MHC Class I & II
Trophoblast 3
Partially SARS-CoV-2 associated
Down-regulation of MHC Class II
Down-regulation of placentitis villous trophoblast associated gene ICAM1

Figure S7

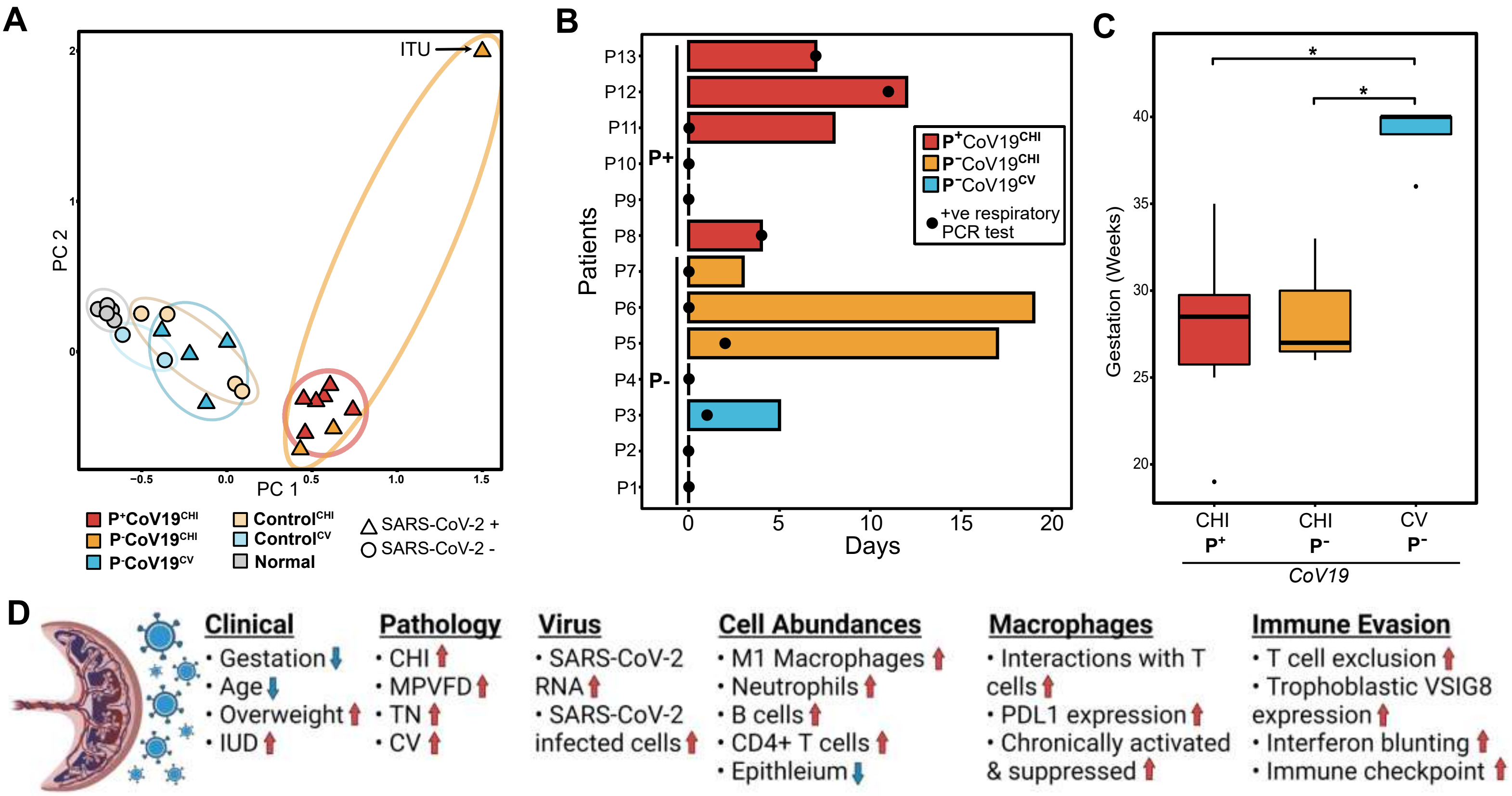

A
ITU
PC 2
PC 1
−0.5
0.0
0.5
1.0
1.5
P+CoV19CHI
P-CoV19CHI
P-CoV19CV
ControlCHI
ControlCV
Normal
SARS-CoV-2 +
SARS-CoV-2 -
B
Patients
P13
P12
P11
P10
P9
P8
P7
P6
P5
P4
P3
P2
P1
P+
P-
Days
0
5
10
15
20
P+CoV19CHI
P-CoV19CHI
P-CoV19CV
+ve respiratory PCR test
C
Gestation (Weeks)
20
25
30
35
40
*
*
CHI P+
CHI P-
CV P-
CoV19
D
Clinical
• Gestation
• Age
• Overweight
• IUD
Pathology
• CHI
• MPVFD
• TN
• CV
Virus
• SARS-CoV-2 RNA
• SARS-CoV-2 infected cells
Cell Abundances
• M1 Macrophages
• Neutrophils
• B cells
• CD4+ T cells
• Epithleium
Macrophages
• Interactions with T cells
• PDL1 expression
• Chronically activated & suppressed
Immune Evasion
• T cell exclusion
• Trophoblastic VSIG8 expression
• Interferon blunting
• Immune checkpoint

| | Maternal respiratory SARS-CoV-2+ | | | | Pre-pandemic controls | | |
|---|---|---|---|---|---|---|---|
| *COVID-19 status* | All | M+P+ | M+P- | M+P- | NA | NA | NA |
| *Pathology* | All | CHI | CHI | CV | CHI | CV | Norm |
| *Sample size* | n=13 | n=6 | n=3 | n=4 | n=4 | n=2 | n=5 |
| Clinical characteristics n (%) | | | | | | | |
| **Age** | | | | | | | |
| 18-25 | 4 | 3 (50) | 1 (33) | 0 (0) | 0 (0) | 0 (0) | 2 (40) |
| 26-35 | 6 | 2 (33) | 1 (33) | 3 (75) | 2 (50) | 1 (50) | 3 (60) |
| 36+ | 3 | 1 (17) | 1 (33) | 1 (25) | 2 (50) | 1 (50) | 0 (0) |
| **Foetal Sex** | | | | | | | |
| Male | 6 | 2 (33) | 2 (67) | 2 (50) | 1 (25)* | 1 (50) | 4 (80) |
| Female | 6 | 4 (67) | 1 (33) | 1 (25) | 2 (50) | 1 (50) | 1 (20) |
| Male/female twin | 1 | 0 (0) | 0 (0) | 1 (25) | 0 (0) | 0 (0) | 0 (0) |
| **Ethnicity** | | | | | | | |
| Caucasian | 5 | 3 (50) | 1 (33) | 1 (1) | 2 (50) | 2 (100) | 3 (60) |
| Black | 2 | 2 (33) | 0 (0) | 0 (0) | 0 (0) | 0 (0) | 1 (20) |
| Asian | 4 | 1 (17) | 1 (33) | 2 (50) | 1 (25) | 0 (0) | 1 (20) |
| Middle eastern | 2 | 0 (0) | 1 (33) | 1 (25) | 1 (25) | 0 (0) | 0 (0) |
| **Body mass index** | | | | | | | |
| Normal | 2 | 1 (17) | 0 (0) | 1 (25) | 1 (25)* | NA* | NA* |
| Overweight | 8 | 4 (67) | 2 (67) | 2 (50) | 0 (0) | NA | NA |
| Obese | 3 | 1 (17) | 1 (33) | 1 (25) | 2 (50) | NA | NA |
| **Gestation** | | | | | | | |
| Second trimester | 3 | 2 (33) | 1 (33) | 0 (0) | 1 (25) | 0 (0) | 0 (0) |
| Third trimester | 10 | 4 (67) | 2 (67) | 4 (100) | 3 (75) | 2 (100) | 5 (100) |
| **COVID-19 Respiratory illness** | | | | | | | |
| Severe | 1 | 0 (0) | 1 (33) | 0 (0) | NA | NA | NA |
| Mild | 7 | 5 (83) | 1 (33) | 1 (25) | NA | NA | NA |
| Asymptomatic | 5 | 1 (17) | 1 (33) | 3 (75) | NA | NA | NA |
| **Obstetric outcome** | | | | | | | |
| Intra uterine death | 8 | 6 (100) | 2 (67) | 0 (0) | 1 (25) | 0 (0) | 0 (0) |
| Prematurity | 2 | 0 (0) | 1 (33) | 1 (25) | 1 (25) | 2 (100) | 0 (0) |
| Intra-uterine growth restriction | 2 | 0 (0) | 1 (33) | 1 (25) | 2 (50) | 1 (50) | 0 (0) |
| Fetal distress | 3 | 0 (0) | 0 (0) | 3 (75) | 0 (0) | 0 (0) | 0 (0) |
| Pathological features n (%) | | | | | | | |
| **Macroscopic findings** | | | | | | | |
| MPVFD | 9 | 6 (100) | 3 (100) | 0 (0) | 2 (50) | 0 (0) | 0 (0) |
| Abruption | 1 | 0 (0) | 1 (33) | 0 (0) | 0 (0) | 1 (50) | 0 (0) |
| Membrane opacity | 7 | 3 (50) | 1 (33) | 3 (75) | 1 (25) | 1 (50) | 0 (0) |
| **Microscopic findings** | | | | | | | |
| Chronic histiocytic Intervillositis | 10 | 6 (100) | 3 (100) | 1 (25)[ǂ] | 4 (100) | 0 (0) | 0 (0) |
| Chronic villitis | 9 | 4 (67) | 3 (67) | 3 (75) | 1 (25) | 2 (100) | 0 (0) |
| Basal villitis | 6 | 2 (33) | 1 (33) | 3 (75) | 1 (25) | 2 (100) | 0 (0) |
| Acute chorioamnionitis | 5 | 1 (17) | 2 (67) | 2 (50) | 0 (0) | 0 (0) | 0 (0) |
| Chronic chorioamnionitis | 7 | 5 (83) | 1 (33) | 1 (25) | 1 (25) | 2 (100) | 0 (0) |
| Chronic deciduitis | 9 | 4 (67) | 2 (67) | 3 (75) | 2 (50) | 2 (100) | 0 (0) |
| Secondary bacterial infection | 1 | 0 (0) | 0 (0) | 1 25) | 0 (0) | 0 (0) | 0 (0) |
| Funisitis | 2 | 0 (0) | 0 (0) | 2 (50) | 0 (0) | 0 (0) | 0 (0) |
| Sub-chorionic microabscesses | 1 | 0 (0) | 1 (33) | 0 (0) | 0 (0) | 0 (0) | 0 (0) |

Supplementary Table 1: Clinicopathological features of cases and controls. Key: [ǂ] One patient showed very focal intervillous histiocytosis; * Data for this variable is incomplete for this subgroup % percentage. Abbreviations: NA = not applicable/available; n = number; MPVFD = massive pervillous fibrin deposition; CHI = chronic histiocytic intervillositis; CV = chronic villitis; norm = normal; M+ = positive maternal SARS-CoV-2 respiratory infection; M- = negative maternal SARS-CoV-2 respiratory infection; P+ = positive SARS-CoV-2 trophoblast infection; P- negative SARS-CoV-2 trophoblast infection.

| Cycle position | Total volume | Mouse antibodies | Concentration (1 in x) | Volume of antibody | Rabbit antibodies | Concentration (1 in x) | Volume of antibody |
|---|---|---|---|---|---|---|---|
| 1 | 450 | CD45RO | 50 | 9 | Nucleocapsid | 2000 | 0.225 |
| 2 | 450 | CD45 | 30 | 15 | IP10 | 50 | 9 |
| 3 | 450 | Mum1 | 20 | 22.5 | CD80 | 200 | 2.25 |
| 4 | 450 | CD15 | 200 | 2.25 | Granzyme | 300 | 1.5 |
| 5 | 450 | Vimentin | 50 | 9 | ACE2 | 50 | 9 |
| 6 | 450 | CD68 | 50 | 9 | FoxP3 | 50 | 9 |
| 7 | 450 | CD8 | 50 | 9 | PD1 | 500 | 0.9 |
| 8 | 450 | Ki67 | 25 | 18 | PDL1 | 50 | 9 |
| 9 | 450 | CK | 50 | 9 | CD4 | 200 | 2.25 |
| 10 | 450 | CD20 | 100 | 4.5 | CD3 | 500 | 0.9 |
| 11 | 450 | Spike | 200 | 2.25 | CD163 | 200 | 2.25 |
| 12 | 450 | NA | NA | NA | Fibrinogen | 3000 | 0.15 |
| 13 | 450 | SMA | 1000 | 0.45 | GPNMB | 250 | 1.8 |
| 14 | 450 | | | | CD39 | 300 | 1.5 |
| 15 | 450 | | | | CD103 | 100 | 4.5 |
| 16 | 450 | | | | CTLA4 | 200 | 2.25 |
| 17 | 450 | | | | CD56 | 200 | 0.5 |

Supplementary Table 2: Antibody pairings, species, volumes, and concentrations for the Lunaphore COMET panel.
